# ACAT: A Fast and Powerful P-value Combination Method for Rare-variant Analysis in Sequencing Studies

**DOI:** 10.1101/482240

**Authors:** Yaowu Liu, Sixing Chen, Zilin Li, Alanna C. Morrison, Eric Boerwinkle, Xihong Lin

## Abstract

Set-based analysis that jointly tests the association of variants in a group has emerged as a popular tool for analyzing rare and low-frequency variants in sequencing studies. The existing set-based tests can suffer significant power loss when only a small proportion of variants are causal, and their powers can be sensitive to the number, effect sizes and effect directions of the causal variants and the choices of weights. Here we propose an Aggregated Cauchy Association Test (ACAT), a general, powerful and computationally efficient p-value combination method to boost power in sequencing studies. First, by combining variant-level p-values, we use ACAT to construct a set-based test (ACAT-V) that is particularly powerful in the presence of only a small number of casual variants in a variant set. Second, by combining different variant set-level p-values, we use ACAT to construct an omnibus test (ACAT-O) that combines the strength of multiple complimentary set-based tests including the burden test, Sequence Kernel Association Test (SKAT) and ACAT-V. Through analysis of extensively simulated data and the whole-genome sequencing data from the Atherosclerosis Risk in Communities (ARIC) study, we demonstrate that ACAT-V complements the SKAT and burden test, and that ACAT-O has a substantially more robust and higher power than the alternative tests.

## INTRODUCTION

With the advent of next generation sequencing technology, whole genome and exome sequencing in large cohorts enables the discovery of low-frequency and rare genetic variation that are likely to have substantial contributions to the “missing heritability” and new genetic discovery of complex traits and diseases^1;2^. For example, an exome sequencing study of human height in > 710,000 individuals identified 83 rare and low-frequency coding variants that explained an additional 1.7% of the height heritability^3^. As rare and low-frequency variants appear infrequently in the population, the standard single-variant analysis that has been applied for common variants in genome-wide association studies (GWAS) is underpowered without very large effect sizes and/or sample sizes^4^. Set-based methods, which jointly analyze variants in a group (e.g., exon variants in a gene), have been proposed and become increasingly popular^4^. These methods perform analysis by grouping rare variants in a set to aggregate their small and moderate effects to increase statistical power.

Over the past a few years, the Sequence Kernel Association Test (SKAT)^5^ and burden tests^6–8^ have emerged as the most widely-used methods for set-based rare-variant analysis, partly because of their undemanding computational requirement, flexibility to adjust covariates for analyzing both binary and quantitative data, and ability to incorporate functional annotations and to allow for related subjects. In order to be power-robust to the directionality of variant effects, an omnibus test SKAT-O^9^ was also proposed to combine SKAT and burden test statistics adaptively based on the observed data. However, SKAT, SKAT-O and burden tests can lose substantial power under sparse alternatives^10^; ^11^, i.e., only a small proportion of variants in a set are associated with a disease/trait. Sparse alternatives are natural and reasonable hypothesis in sequencing studies, as most variants in a set are anticipated to have no influence on the risk or related traits of a disease. The exponential combination test^12^ was proposed to improve power in the sparse situation, but it requires permutation to evaluate the set significance, which is computationally burdensome or even infeasible for large-scale whole genome sequencing studies.

The power of different set-based tests depends on the underlying genetic architecture, which may differ in the numbers, effect sizes and effect directions of the casual variants in different variant sets. For instance, a proper choice of weights in SKAT and burden tests can boost the power substantively for rare-variant analysis. Wu et al.^5^ proposed to use the family of beta densities of minor allele frequencies of the variants in a region as the weights. If rarer variants are more likely to have larger effects, upweighting the rarer variants would enhance the analysis power. However, if all the variants have the same or similar effect sizes, the use of equal weights might be better. In practice, the genetic architecture of complex traits is rarely known in advance and likely to vary from one region to another across the genome and from one trait to another. Another important limitation of the existing set-based tests is that they could suffer a substantial loss of power if their assumptions are violated. Hence, it is desirable to have an omnibus test that combines the strength of multiple tests and is robust to the sparsity of casual variants, directionality of effects, and the choice of weights.

A widely adopted approach for combining multiple tests is to take the minimum p-value of tests as a summary of the significance. This approach, however, often requires numerical simulations to evaluate the significance of the omnibus test and is computationally expensive as the multiple tests are often correlated. We note that SKAT-O also uses the minimum p-value approach to combine SKAT and the burden test, and its p-value can be calculated efficiently without simulations. However, the particular technique of p-value calculation for SKAT-O is not applicable to the combination of different tests in general (e.g., the combination of SKAT tests under different choices of weights). Fisher’s method^13^ can also be used for the combination of complementary tests^14^. However, it suffers from the same computational issue as the minimum p-value method and could result in a considerable loss of power as the combined test statistics are calculated from analyzing the same data and often highly correlated.

In this paper, we propose an Aggregated Cauchy Association Test (ACAT), a flexible and computationally efficient p-value combination method to boost power in sequencing studies. ACAT first transforms p-values to be Cauchy variables, takes the weighted summation of them as the test statistic and then evaluates the significance analytically. ACAT is a general method for combining p-values and can be used in different ways depending on the types of p-values being combined. When applied to combining variant-level p-values, ACAT is a set-based test that is particularly powerful in the presence of a small number of causal variants in a variant set, and therefore complements the existing SKAT and burden test. When applied to combining set-level p-values from multiple variant set tests, ACAT is an extremely fast omnibus testing procedure that performs the multiple testing adjustment analytically and is applicable to the combination of any tests.

The most distinctive feature of ACAT is that it only takes the p-values (and weights) as input and the p-value of ACAT can be well approximated by a Cauchy distribution. Specifically, neither the linkage disequilibrium (LD) information in a region of the genome nor the correlation structure of set-level test statistics are needed for calculating the p-value of ACAT. This feature offers several advantages. First, the computation of ACAT only involves simple analytic formulae and is extremely fast. Given the variant-level or set-level p-values, applying ACAT for analysis at the whole-genome scale requires just a few seconds on a single laptop. Second, as a set-based test, ACAT only requires variant-level summary statistics (from a single study or meta-analysis) and no population reference panel is needed. Third, when the p-values aggregated by ACAT are calculated from appropriate models that correct for spurious association due to cryptic relatedness and/or population stratification, then ACAT also automatically controls for the spurious association. Another important feature of ACAT is that it allows flexible weights that can be used to incorporate prior information such as functional annotations to further boost power.

For analyzing rare and low-frequency variants, we adapt ACAT to construct set-based tests to increase the analysis power in sequencing studies. We first propose a set-based test ACAT-V, which combines the variant-level p-values and has strong power against sparse alternatives. As mentioned earlier, SKAT and burden tests have limited power if most variants in a set are not associated with the trait. In contrast, the proposed ACAT-V could also lose power in the presence of many weakly associated variants. In addition, the choice of weights could also have a substantial impact on the analysis power. Therefore, we further propose to combine the evidence of association from SKAT, the burden test and ACAT-V, each with two types of weights (i.e., equal weights and weights that upweight rare variants), to improve the overall power. We use ACAT to combine the p-values of the multiple set-based tests and refer to this omnibus test as ACAT-O.

We conducted extensive simulations to investigate the type I error of ACAT-V and ACAT-O and compare their power with that of alternative set-based tests across a broad range of genetic models for both continuous and dichotomous traits. Through the analysis of Atherosclerosis Risk in Communities (ARIC) whole-genome sequencing data^15^, we demonstrate the complementary performance of ACAT-V, SKAT and the burden test, and that ACAT-O identifies more significant regions than each individual test and is very robust across different studies. A summary of the proposed methods and their relationships are provided in **Figure 1**.

**Figure 1.**
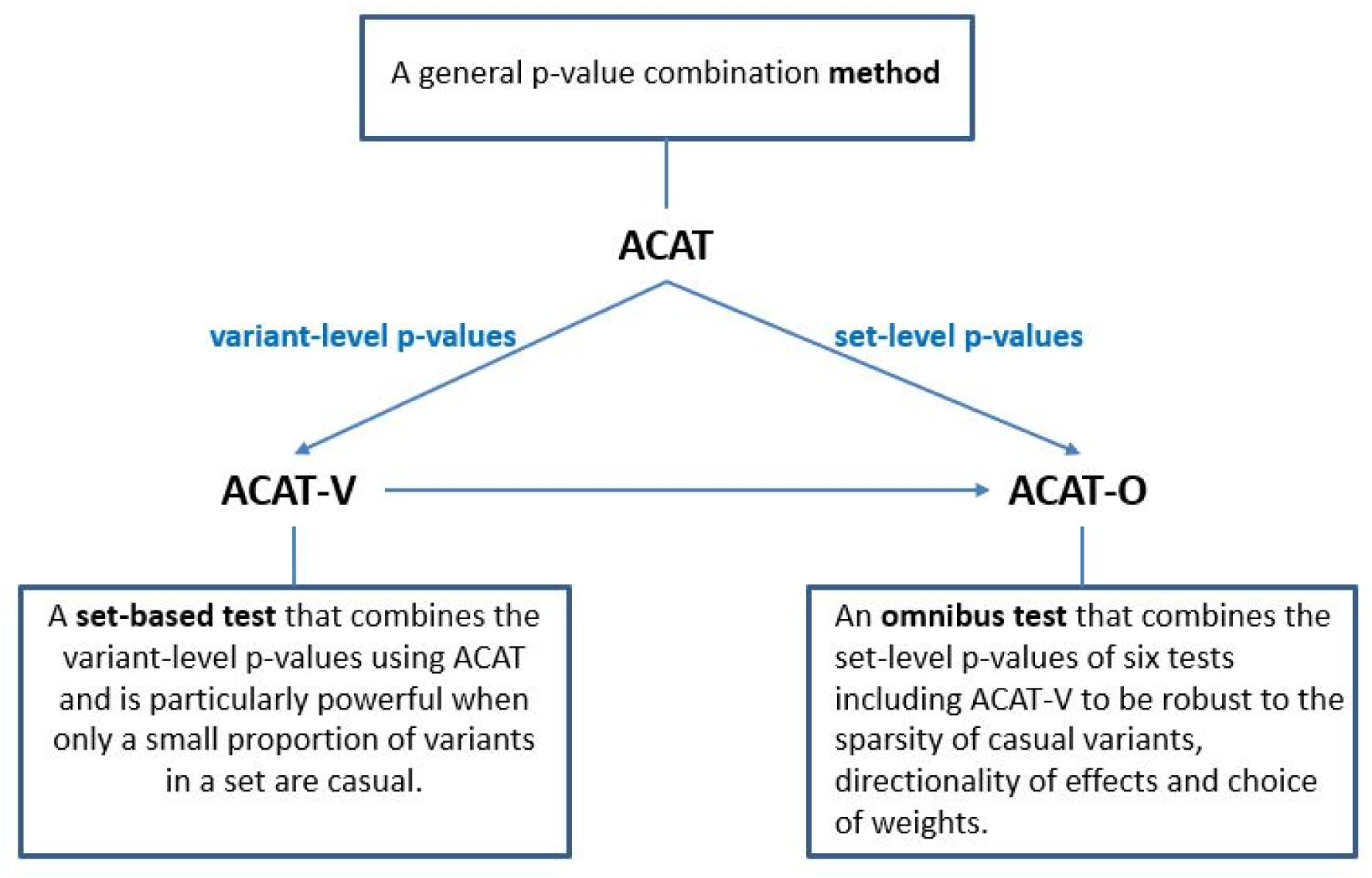
Summary of the proposed methods ACAT, ACAT-V and ACAT-O, and the relationship among them.

## MATERIAL AND METHODS

### Aggregated Cauchy Association Test

The aggregated Cauchy association test (ACAT) is a general and flexible method to combine p-values, which can represent the statistical significance of different kinds of genetic variations in sequencing studies. Let *p*_1_, *p*_2_, …, *pk* denote the p-values combined by ACAT. For set-based testing, the p-values may correspond to *k* variants in a region and can be calculated from a variety of models for single-variant analysis, including linear regression or mixed model for continuous traits and logistic regression or mixed model for binary traits. For the combination of different tests, *p*_*i*_’s can be the p-values of *k* different set-based tests (such as SKAT, the burden test or tests with different choices of weights) for a same region. As ACAT only aggregates p-values, cryptic relatedness and/or population stratification are automatically controlled by fitting appropriate models that *p*_*i*_’s are calculated from through methods such as principal components analysis^16^ or mixed models^17^; ^18^.

Similar to the classical Fisher’s test^13^, ACAT uses a linear combination of transformed p-values as the test statistic, except that the p-values are transformed to follow a standard Cauchy distribution under the null hypothesis and flexible weights are allowed in the combination. Specifically, the ACAT test statistic is

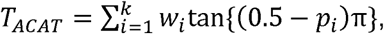

where *p*_*i*_’s are the p-values, *w*_*i*_’s are non negative weights, and the transformation tan{(0.5 – *p*_*i*_)*π* } is Cauchy distributed if *p*_*i*_ is from the null.

For both set-based analysis and combination of tests, the p-values are expected to have moderate or strong correlations due to the LD of different variants or the fact that the p-values represent the significance of different tests of a same region based on the same data. The most distinctive feature of ACAT is that the null distribution of the test statistic *TACAT* can be well approximated by a Cauchy distribution with a location parameter 0 and a scale parameter 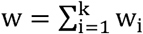 without the need to estimate and account for the correlation among p-values^19^. Therefore, based on the cumulative density function of the Cauchy distribution, the p-value of ACAT is approximated by

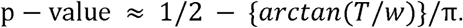

Because this approximation does not need information about the correlation of p-values, calculating the p-value of ACAT requires almost negligible computation and is extremely fast. For instance, given the summary statistics of variants, set-based analysis using ACAT for the whole genome can be performed on a single laptop and only takes a few seconds. Furthermore, the approximation is particularly accurate when ACAT has a very small p-value, which is a very useful feature in sequencing studies as only very small p-values can pass the stringent genome-wide significance threshold and are of particular interest. The reason that ACAT maintains these notable features is due to the heavy tail of the Cauchy distribution, which makes the distribution of the test statistic *TACAT* (especially the tail of the distribution) insensitive to the correlation of the p-values and can be approximated without the correlation information. See Appendix A for more details about the theoretical justification of the Cauchy-distribution-based approximation and general practical guidelines regarding its approximation accuracy.

### ACAT-V for rare-variant association analysis

We use ACAT to combine variant-level p-values and develop a set-based test (ACAT-V) that is particularly powerful under sparse alternatives, i.e. the presence of a small number of causal variants in a set. The variant-level p-values are calculated based on the normal approximation, which becomes inaccurate as the number of minor alleles decreases. In addition, the behavior of the variant-level p-value/test statistic for a very rare variant under the alternative will be attenuated towards the null in comparison to a common variant for a given effect size due to the extremely low number of minor alleles. Therefore, a direct application of ACAT to aggregate the variant-level p-values in a region would result in an overly conservative type I error and lowered power. To address this issue, we first use the burden test to aggregate variants with a minor allele count (MAC) less than a certain number, e.g., 10, and then combine the p-value of this burden test of very rare variants with variant-level p-values of the other variants in a region using ACAT. Specifically, let *p*_0_ denote the p-value of the burden test and *p*^1^*,…,ps* denote the variant-level p-values of variants with a MAC ≥ 10. The ACAT-V test statistic is

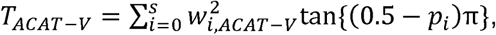

where 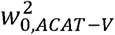 is the weight for the burden test p-value of variants with MAC<10 and 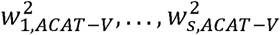 are weights for the other individual variants. For dichotomous traits, the normal approximation could be inaccurate even for variants with MAC ≥ 10 and we use a saddlepoint approximation^20^ to improve the accuracy of the variant-level p-values.

Similar to SKAT and the burden test, including appropriate weights in ACAT-V can yield improved power. One natural and straightforward way of choosing weights for ACAT-V is to use the same weights as in SKAT. However, the weights in SKAT and ACAT-V are not directly comparable and have different interpretations as the SKAT weights are on the individual variant score statistic scale while the ACAT-V weights are on the individual variant transformed p-value scale.

Specifically, the SKAT test statistic^4^ can be written as

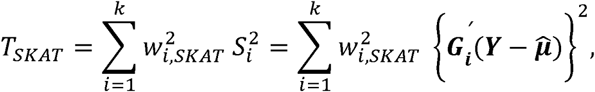

where 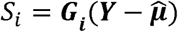 is the single-variant score test statistic, 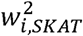 is the weight and ***G***_*i*_ is a vector of allele counts for the *;i*th variant, ***Y*** is the phenotype vector and 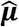 is the predicted mean of, ***Y*** under the null hypothesis. SKAT puts weights on the single-variant score test statistics, whose scales (i.e., variances) are different and depend on the minor allele frequencies (MAF) of variants. In contrast, ACAT-V puts weights on the transformed p-values, which have the same scale and follow a standard Cauchy distribution under the null hypothesis. To make a connection between the weights in SKAT and ACAT-V, we can standardize the single-variant score test statistics to have the same scale by multiplying the inverse of the sample standard derivations of the variants 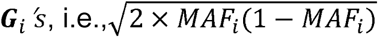. Hence, we can set the weights in ACAT-V as

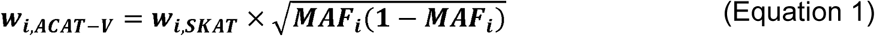

to make them comparable with those in SKAT.

For rare-variant analysis, Wu et al.^5^ proposed to set *w*_*i,SKAT*_= *Beta*(*MAF*_*i*_,*a*_1_,*a*_2_), the beta density function with two parameters *a*_1_and *a*_2_ evaluated at the MAF of the *i*-th variant in a region. Common choices of the parameters are *a*_1_ *=1* and *a*_2_*= 25*, which correspond to the assumption that rarer variants have larger per-allele effect, and *a*^1^*= a*^2^= 1, which corresponds to the assumption of same per-allele effect for all variants. As a demonstrating example, we will also use the beta densities and Equation (1) to set weights for ACAT-V in our simulations and real-data analyses.

Because the p-value *p*_0_(and *MAF*_0_) of the burden test for variants with MAC<10 in ACAT-V corresponds to multiple variants, we set *MAF*_0_ to be the average MAF of the variants with MAC< 10 and calculate the weight (*w*^0^*, ACAT-V*) for the burden test p-value based on Equation (1) accordingly. In addition, the weights in this burden test for variants with MAF<10 also needs to be specified. In fact, there are two layers of weights in ACAT-V. Here we choose the two layers of weights to have the same type to be consistent. For instance, if the beta density *Beta*(1,25) is applied for weighting in ACAT-V, it is used to weight the p-values in the outer layer based on Equation (1) and also used to weight the variants with MAC< 10 in the burden test in the inner layer.

While we choose weights based on the MAFs of the variants here, other forms of weights, such as those based on the functional annotations, can also be used. As long as the weights do not depend on the phenotypes, the p-value of ACAT-V can be approximated efficiently via the Cauchy distribution.

### An omnibus test: ACAT-O

In the ACAT-V test statistic, the Cauchy transformed p-values increases very fast as the p-value approaches to 0 and the weighted summation is essentially dominated by the components with very small p-values. Therefore, ACAT-V mainly uses a few smallest p-values to represent the significance of a region and is particularly powerful when only a small number of variants are casual. In contrast, the SKAT and burden test are more powerful than ACAT-V if a region contains a moderate or large number of casual variants. Further, in the case of a high proportion of causal variants, the burden test could have stronger power than SKAT if the casual variants have the same direction of association but lose power compared with SKAT if the effects of casual variants are bi-directional. The choice of weights could also affect the power of a set-based test. If the effect size of a causal variant has a negative relationship with the MAF of a variant, then the beta weights using MAF with parameters α _1_=1 and α _2_= 25 would lead to stronger power than the weights with α _1_= α _2_=1 (i.e., the equal weights). But if the negative relationship is not true (for instance, there is no relationship between the effect size and MAF), the weights with α _1_= α _2_=1 may be better. In practice, we rarely have the prior information about the number, effect sizes and effect directions of underlying casual variants, which could also vary from one region to another across the genome and from one trait to another. Therefore, it is desirable to determine the test adaptively based on the observed data to combine the strength of multiple tests. We can use ACAT to combine the p-values of multiple set-based tests to construct an omnibus test (ACAT-O) that has robust power under various genetic architectures. While the p-values of set-based tests correspond to the significance of a same region and are (highly) correlated, ACAT does not require the correlation structure of the p-values of different set-based tests, and therefore is well-suited for omnibus testing.

We construct the omnibus test (ACAT-O) for a variant set by combining six set-based tests, i.e., ACAT-V, SKAT, burden test and each test with two choices of weights (i.e., the weights with α _1_= α _2_*=1* and weights with α _1_=1 and α _2_= 25), to be robust to the sparsity of causal variants, directionality of effects, and the choice of weights. Specifically, the ACAT-O test statistic is

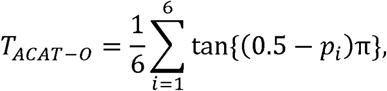

where p_i_’s, are the p-values of the six tests and the tests are treated equally in the combination. We will apply this test in all the simulations and real data analyses. One can also use ACAT to combine other set-based tests. The p-value of ACAT-O can be calculated very fast via the Cauchy-distribution-based approximation.

As the underlying true genetic architecture is seldomly known in advance, it is possible that some tests incorporated by ACAT-O do not have sufficient statistical power and therefore would lead to loss of power in the omnibus testing. For example, if the protective and harmful variants in a region have the same numbers and effect sizes, the burden test would be powerless. Thus, it is desirable that the omnibus testing procedure is not sensitive to the inclusion of underpowered tests such that the power loss can be minimized and does not exceed the power gain from other powerful tests in the omnibus testing. As mentioned earlier, ACAT mainly focuses on the few smallest p-values, which is also an attractive feature for omnibus testing and makes ACAT-O robust even when some of the incorporated tests are underpowered. The minimum p-value method also has a similar feature. However, an advantage of ACAT over the minimum p-value method is that calculation of the p-value of ACAT does not require accounting for the correlation of the individual tests, while the minimum p-value method requires estimating and accounting for the correlation of the individual tests, which is often difficult and time-consuming.

### Simulation Studies

We carried out extensive simulations to investigate the type I error of ACAT-V and the omnibus test ACAT-O and compare their power with SKAT and the burden test under different choices of weights. For all the simulations, we generated 100 1Mb regions of sequencing genotype data based on a calibration coalescent model that mimics the LD structure and local recombination rate of the European population^21^. Our simulation studies focus on rare and low-frequency variants, so we excluded variants with MAF>0.05 in all of the 1 Mb regions.

#### Simulations of Type I error

To obtain a total of 10_8_phenotype-genotype data sets, we first randomly selected 1000 4 kb sub-regions from each of the 100 1Mb regions and then generated 1000 phenotypes for each 4 kb sub-region of genotype data. The variant set length of 4kb is from a sliding window approach^15^ and will also be used in the real-data analysis described in the next section. As it is common to adjust for covariates such as age, gender and principal components in practice, we included four associated covariates (three continuous and one binary) in the null model for both continuous and dichotomous traits. Specifically, we simulated continuous phenotypes according to the linear model:

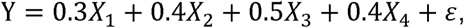

and dichotomous phenotypes according to the logistic model:

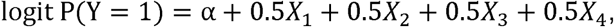

where *x*_1_,*x*_2_,*x*_3_are generated independently from a standard normal distribution, *x*_4_takes values 0 and 1 with equal probability,αis an error term following a standard normal distribution, α was determined to have a prevalence of 0.01 and balanced case-control sampling is used for dichotomous trait. We set the sample size *n* to be 2500, 5000, 7500 and 10000. For each test, the empirical type I error rate is calculated as the proportion of p-values less than the significance level.

#### Simulations of Empirical Power

To assess the power performance of competing set-based tests, we randomly selected causal variants within each of the 4kb regions to simulate phenotypes under the alternative. Specifically, we generated continuous phenotypes by

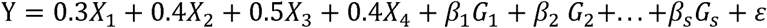

and dichotomous phenotypes by

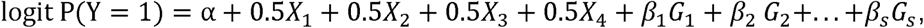

where G_1_,G_2_ …G_3_ are the genotypes of randomly selected casual variants, βi’s are the effect sizes for the casual variants and the other symbols are as defined for the simulations of type I error.

We varied the proportion, effect sizes and effect directions of casual variants to investigate the impact of these three factors on the power of different tests. The proportion of casual variants was set to be 5%, 20% and 50% to cover the situations of sparse and dense signals. The casual variants in a region could be all deleterious or protective, or some of them are protective and others deleterious. Hence, we examined two settings of effect directions: the signs of βi’s are either in the same direction or are determined randomly and independently with an equal probability of 0.5. We also investigated two scenarios of effect sizes: *β_i_*’s, have the same magnitude *b* or are set to be *c*|log^10^ *MAF*_*i*_ *|* such that variants with smaller MAF have larger effects, where constants *b* and *c* depend on the proportions of casual variants and their values are presented in **Table S1**. We considered all possible combinations of the three factors (i.e., the proportion, sizes and directions of nonzero *β_i_*’s,) and had 12 simulation configurations in total that covered a variety of genetic architecture scenarios. The significance level *α* was set to be 10^−6^ to mimic genome-wide studies and the empirical power of each test was estimated as the proportion of p-values less than *α* on the basis of 10^3^ replicates.

### ARIC whole genome sequencing data

The Atherosclerosis Risk in Communities (ARIC) study has been described in detail previously^22^. Regarding the whole genome sequencing data, DNA samples were sequenced at 7.8-fold average depth on Illumina HiSeq instruments and genotyping was performed at the Baylor College of Medicine Human Genome Sequencing Center. After sample-level quality control detailed in Morrison et al^15^, there were around 55 million variants in 1,860 African American (AA) participants and 33 million variants in 1705 European American (EA) participants. Among all the variants, 17.3% and 19.4% of them are common variants (MAF>5%) in the AA and EA populations, respectively. Our study primarily focuses on analyzing low-frequency (1%≤MAF≤ 5%, 13.4% in AA and 9.1% in EA) and rare (MAF < 1%, 69.3% in AA and 71.5% in EA) variants across the genome using a sliding window approach^15^ that chooses physical windows of 4kb in length as the analytical units and starts at position 0 bp for each chromosome with a skip length of 2kb. A minimum number of 3 minor allele counts is required in a window, resulting in a total of 1,337,673 and 1,337,382 4kb overlapping windows in AA and EA, respectively. The distribution of the number of variants in a window has a median of 60 in AA and a median of 37 in EA, and is highly skewed to the right.

The example application presented here focuses on the analysis of two quantitative traits (lipoprotein(a) (Lp(a)) and neutrophil count). The methods for the measurement of each trait were described in detail in Morrison et al^15^. We adjusted for age, sex and the first three principal components for both traits and additionally included current smoking status as a covariate in the analysis of neutrophil count. Because the distributions of both Lp(a) and neutrophil count are markedly skewed, we applied rank-based inverse normal transformation^23^ to both traits and used the transformed traits as phenotypes in the analyses. For each 4kb window, we performed set-based association using ACAT-O, ACAT-V, SKAT and burden test, and weighting the variants by beta weights using MAFs with parameters either α_1_ = α_2_ =1 or α_1_ =1 and α^2^ = 25. As around 1.3 million windows are tested in each analysis, we used the Bonferroni method and set the genome-wide significance threshold at 3.75×10^−8^ (approximately equal to 0.05/1,337,000).

## RESULTS

### Simulation of the Type I Error

The empirical type I error rates for ACAT-V and ACAT-O are presented in **Table 1** for significance levels α= 10^−4^,10^−5^, and 10^−6^. The results demonstrate that the type I error rate is protected for both continuous and dichotomous traits, although slightly conservative for very small significance levels (e.g., α= 10^−6^). We note that the conservativeness is not due to the Cauchy-distribution-based approximation, but the conservativeness of the p-values of the tests that are aggregated by ACAT. In fact, the theory provided in Appendix A suggests that the Cauchy-distribution-based approximation becomes more accurate as the significance level decreases when the aggregated p-values follow a uniform distribution between 0 and 1 exactly. However, the p-values of set-based tests (e.g., SKAT) or the variant-level p-values are conservative for rare variants and dichotomous traits^5^, which results in the slightly deflated type I error rate of ACAT-V and ACAT-O.

**Table 1.**
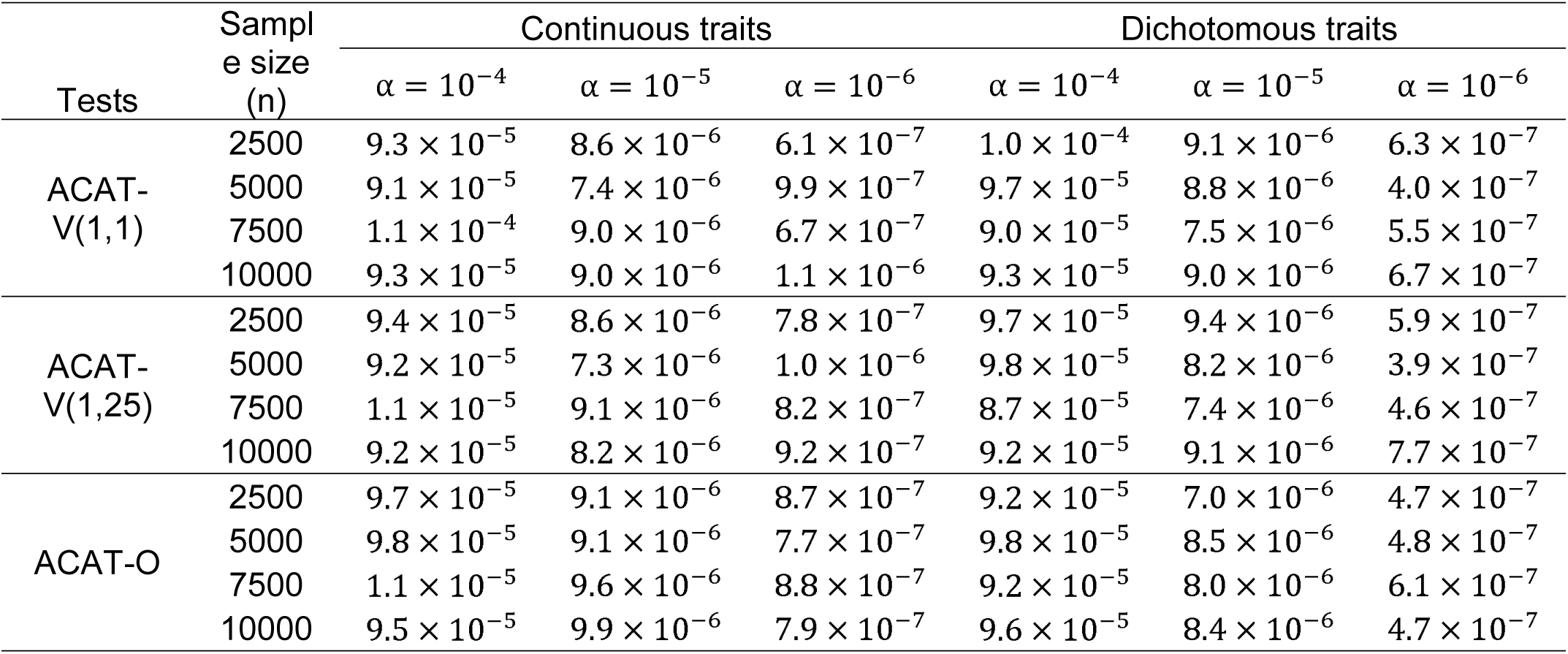
Type I error estimates of ACAT-V and ACAT-O aimed at testing the association between randomly selected 4kb regions with a continuous or dichotomous trait. In each 4kb region, common variants with MAF>5% are excluded to focus on only rare and low-frequency variants. Each cell represents type I error rate estimates as the proportion of p values less than α under the null hypothesis on the basis of 10^8^ replicates. For ACAT-V, the two numbers in the parentheses correspond to the beta(MAF) weight parameters α_1_ and α_2_, respectively.

### Simulation of the power

We compared the power of ACAT-V and ACAT-O with SKAT and the burden test under a variety of scenarios for both continuous and dichotomous traits. Except for the omnibus test ACAT-O, we considered two choices of weights using MAF for the other three tests, i.e., the beta weights of MAFs with α_1_ = α_2_ =1 and the beta weights of MAF with α_1_=1 and α_2_= 25.**Figures 2 and 3** display the results under the 12 simulation configurations for continuous and dichotomous traits, respectively. When only a small proportion (5%) of variants are casual, ACAT-V had much higher power than SKAT and the burden test regardless of the effect directions and choices of weights. This is expected because ACAT-V mainly uses a few smallest p-values to represent the significance of a region, while SKAT and burden test use a linear combination of the (squared) score statistics and the overall signal strength is diluted by the dominating large number of non-casual variants. ACAT-V still outperformed SKAT and burden test in the presence of a moderate number of casual variants (20%), but had substantial power loss when there are a large proportion of casual variants (50%).

**Figure 2.**
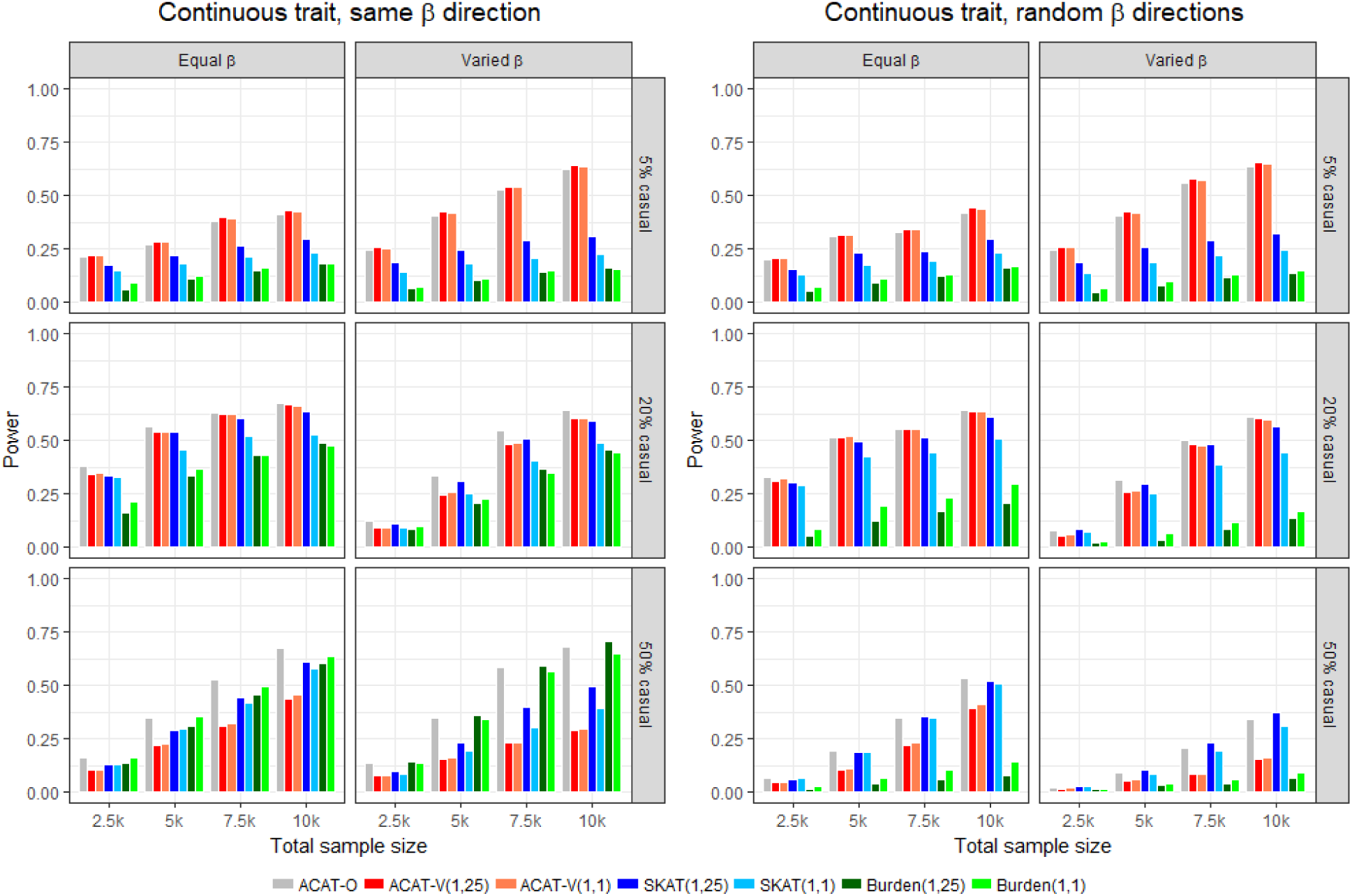
Power comparisons of ACAT-O, ACAT-V, SKAT and burden tests for continuous traits. Each bar represents the empirical power estimated as the proportion of p-values less than α= 10^−6^. The proportion of casual variants is set to be 5%, 20% and 50%, which correspond to the three rows of each panel. The left panel assumes the effects of the causal variants to have the same direction, while the right panel assumes the effect directions are randomly determined with an equal probability. The effect sizes (*|(*β*i |*’*s*) of the casual variants are either all the same as *|(*β*i |=*b (left column in each panel) or have a decreasing relationship with MAF (right column in each panel) as *|(*β_*i*_*|= c*|log^10^ *MAF |*, where constants *b* and *c* depend on the proportions of casual variants and their values are presented in **Table S1**. For each configuration, the total sample sizes considered are 2500, 5000, 7500, and 10000. Seven methods are compared: ACAT-V(1,25), ACAT-V(1,1), SKAT(1,25), SKAT(1,1), Burden(1,25), Burden(1,1) and the omnibus test ACAT-O that combines the other six tests, where the two numbers in the parentheses indicate the choice of the beta(MAF) weight parameters α_1_and α_2_ in the test.

**Figure 3.**
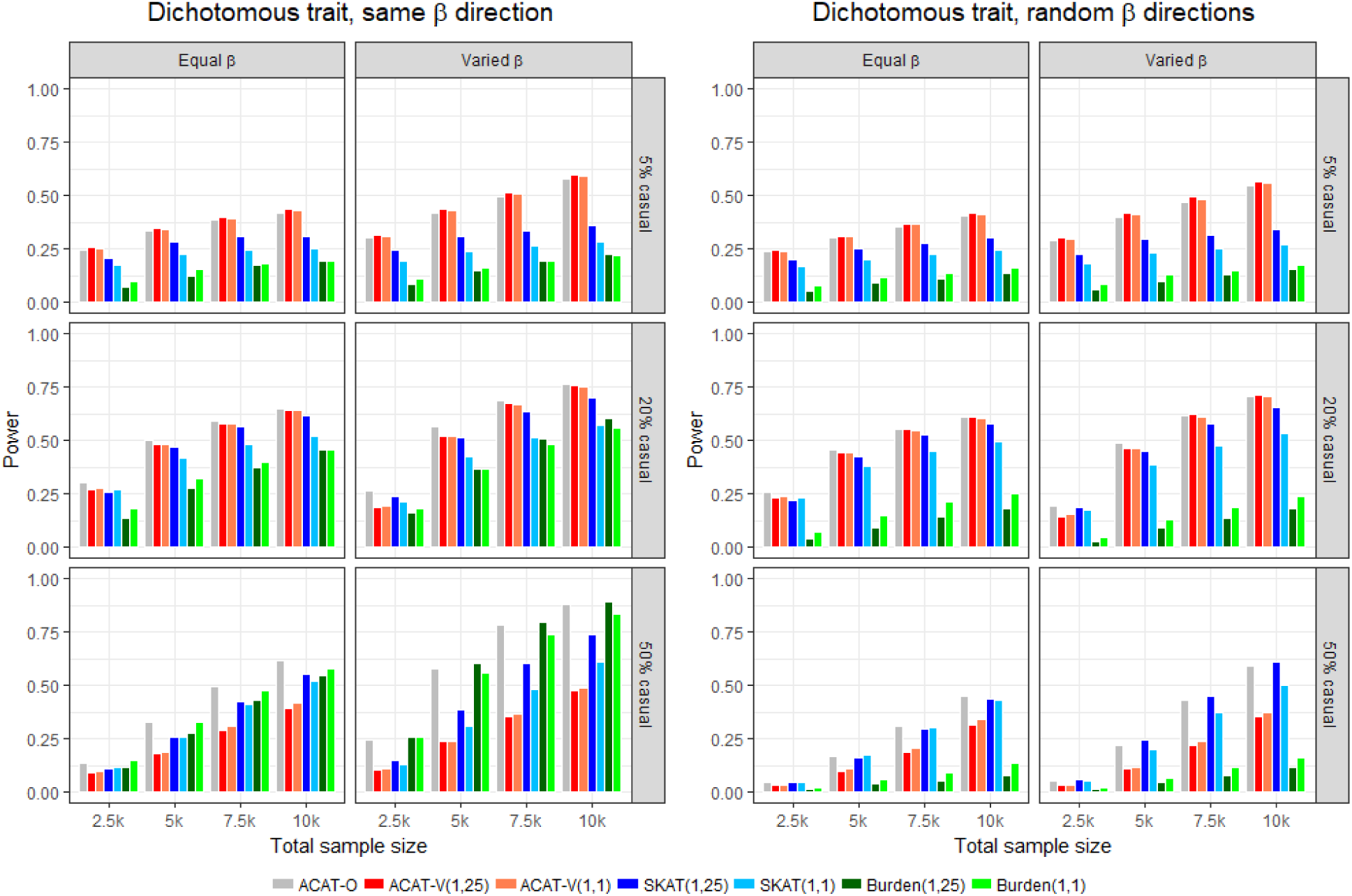
Power comparisons of ACAT-O, ACAT-V, SKAT and burden tests for dichotomous traits. Each bar represents the empirical power estimated as the proportion of p-values less than α= 10^−6^. The proportion of casual variants is set to be 5%, 20% and 50%, which correspond to the three rows of each panel. The left panel assumes the effects of the causal variants to have the same direction, while the right panel assumes the effect directions are randomly determined with an equal probability. The effect sizes (*|*(β_*i*_ ‘*|*’*s*) of the casual variants are either all the same as *|(*β_*i*_ *|*= b (left column in each panel) or have a decreasing relationship with MAF (right column in each panel) as *|(β_i_|= c|*log^1O^ *MAF|*, where constants *b* and *c* depend on the proportions of casual variants and their values are presented in **Table S1**. For each configuration, the total sample sizes considered are 2500, 5000, 7500, and 10000. Seven methods are compared: ACAT-V(1,25), ACAT-V(1,1), SKAT(1,25), SKAT(1,1), Burden(1,25), Burden(1,1) and the omnibus test ACAT-O that combines the other six tests, where the two numbers in the parentheses indicate the choice of the beta(MAF) weight parameters α_1_ and α_2_ in the test.

The burden test was much more sensitive to the effect direction than SKAT and ACAT-V and suffered severe loss of power in the presence of both protective and harmful variants. When a large number of casual variants were present, SKAT exhibited significantly higher power than ACAT-V and burden test in the case of bidirectional effects, while the burden test was more powerful than SKAT and ACAT-V in the case of unidirectional effects with a large number of causal variants. However, all three tests (i.e., ACAT-V, SKAT and burden test) were not robust and could be particularly powerful in some situations but lose substantial power in other situations. In contrast, the omnibus test ACAT-O combined the strength of all the other tests and was very robust to various genetic architectures while losing little power compared to the most powerful test. Indeed, across all the configurations, the power of ACAT-O is either the highest among all the competing methods or just slightly lower than the highest one. In the absence of prior knowledge about the underlying genetic architecture, we expect that ACAT-O can improve the overall power and yield more significant findings than the other methods.

### Application to the ARIC whole genome sequencing data

We applied the proposed methods to analysis of ARIC WGS data. **Table 2** shows the number of 4kb sliding windows identified as significant by each method for Lp(a) and neutrophil count in AA and EA individuals. The significant 4kb sliding windows are also reported in **Table S2, S3 and S4**. None of the set-based tests (SKAT, ACAT-V, or the burden test) under a particular choice of weights consistently exhibited higher statistical power than the other methods across all the analyses. For instance, SKAT(1,25) (the two numbers in the parentheses are the values of beta(MAF) weight parameters α_1_ and α_2_, respectively) detected more windows than SKAT(1,1) in the analyses of Lp(a) in both AAs and EAs, but only detected about half of the significant windows identified by SKAT(1,1) in the analysis of neutrophil count. The relative performance difference between the two traits may suggest that the relationship between the effect sizes and MAFs of variants are trait-specific. Compared to SKAT, the proposed test ACAT-V demonstrated a slightly higher overall power in the analyses of Lp(a) among EAs and was less sensitive to the choice of weights than SKAT. Moreover, **Figures S4-S6** display the scatterplots of the p-values of the significant windows and show that ACAT-V can identify variant sets that are challenging for the SKAT and burden tests to identify.

**Table 2.** The number of significant sliding windows identified by each test (ACAT-O, ACAT-V, SKAT, Burden) that are associated with lipoprotein(a) or neutrophil count among AAs or EAs in the analysis of the ARIC whole genome sequencing data. For ACAT-V, SKAT and the burden test, the two numbers in the parentheses correspond to the beta(MAF) weight parameters α_1_ and α_2_, respectively. The significance threshold α is 3.75x 10^−8^.

| Tests | Lipoprotein(a) |  | Neutrophil count |  |
| --- | --- | --- | --- | --- |
|  | AA | EA | AA | EA |
| ACAT-O | **153** | **46** | **113** | 0 |
| ACAT-V(1,1) | 123 | 42 | 74 | 0 |
| ACAT-V(1,25) | 122 | 41 | 57 | 0 |
| SKAT(1,1) | 103 | 31 | **113** | 0 |
| SKAT(1,25) | 127 | 35 | 62 | 0 |
| Burden(1,1) | 8 | 5 | 20 | 0 |
| Burden(1,25) | 6 | 3 | 8 | 0 |

The burden tests detected markedly fewer significant windows than the other methods, which may result from the bidirectional variant effects on the two traits and/or sparsity of casual variant in most regions. In contrast, the omnibus test ACAT-O was robust across all the analyses. It identified considerably more Lp(a)-associated windows than all the other individual test in both AAs and EAs, and also had a comparable performance to SKAT(1,1) in the analysis of neutrophil count. These results are consistent with our simulation studies, in which ACAT-O either achieved or are very close to the highest power among all the methods and are very robust to diverse genetic architectures.

To facilitate further insights into the performance of different tests, we also presented the genomic landscapes of the windows that were significantly associated with Lp(a) among AAs, Lp(a) among EAs and neutrophil count among AAs in **Figure S1, S2 and S3**, respectively. Overall, the results of SKAT and ACAT-V complemented each other, indicating that both situations of dense and sparse causal variants could appear in different regions across the genome. By combining the complementary results, ACAT-O covered the majority of windows identified by each method and achieved substantial power gain compared to the individual tests. For the Lp(a) trait, the significant windows reside in an 850 kb region on chromosome 6 that includes five genes (*PLG*, *SLC22A2*, *SLC22A3*, *LPA* and *LPAL2*). Previous studies have also identified common variants in these five genes that are significantly associated with Lp(a). Two common variants in *LPA*, which encodes the apolipoprotein(a) component of the Lp(a) lipoprotein particle, showed very strong association and explained 36% of the variation of Lp(a) level^24^. Several intronic variants of *LPAL2* and *PLG* were also found to be strongly associated with Lp(a)^25^; ^26^. The *SLC22A3-LPAL2-LPA* gene cluster has been identified as a strong susceptibility locus for coronary artery disease^27^, which an increased level of the Lp(a) lipoprotein is an independent risk factor for.

For neutrophil count, all of the significant sliding windows reside in a 7.2 Mb region on chromosome 1. SKAT(1,1) was the most powerful approach in the analysis of neutrophil count, but SKAT(1,1) did not identify any significant association with Lp(a) in *LPAL2* among EA individuals or in *PLG* among EA and AA individuals. This illustrates that the genetic architecture varies across different regions and traits, and a single test such as SKAT(1,1) is not robust and can miss important regions in some analyses. ACAT-V detected some unique regions and complemented the SKAT and burden test. For instance, ACAT-V had a wider significant area defined by significant windows surrounding *LPAL2* than SKAT and the results are consistent in both AA and EA populations, suggesting that casual variants might sparsely spread over the region surrounding this gene. Many variants in these unique regions identified by ACAT-V also have large CADD^28^ Phred scores (**Figures S7-S9**), which indicates that these regions are likely to contain functional variants. The omnibus test ACAT-O was able to detect the majority of windows that are only significant by ACAT-V or SKAT and thus had the most robust performance across all the analyses. While the burden tests were substantially less powerful than ACAT-V and SKAT in our analyses, ACAT-O only suffered little loss of power and was also robust to the incorporation of underpowered tests. Hence, ACAT-O not only enables the identification of more significant findings, but also is less likely to miss important regions.

## DISCUSSION

We have proposed ACAT as a general and flexible method for combining p-values and used ACAT to develop two set-based tests (ACAT-V and ACAT-O) for association analysis in sequencing studies. Through extensive simulation studies and analysis of the ARIC whole genome sequencing data using a sliding window approach, we demonstrated that ACAT-V is a powerful test to complement the SKAT and burden test in the presence of a small number of casual variants in a set, and that power improvement can be achieved by combining the p-values of multiple complementary tests using ACAT-O. Our simulations also show that the type I error rates of ACAT-V and ACAT-O are protected for both continuous and dichotomous traits although slightly conservative for very small significance levels.

The most important feature of ACAT is that its p-value can be accurately approximated without the need to account for the correlation of p-values of individual tests, which makes the computation extremely fast. This remarkable feature also enables a wide range of applications of ACAT to various genetic studies beyond the rare-variant analysis considered in this paper.

When used to combine variant-level p-values, ACAT can also be applied to analyses of pathways, genes, gene-sets, gene-environment interactions, or common variants in genomewide association studies (GWAS). In these analyses, ACAT requires only summary statistics (or p-values) instead of individual-level data to test the association between a trait and a group of genetic variants. Analyses of summary statistics protects privacy by circumventing the need for sharing individual-level data and offers huge computational advantages. In addition, compared to other methods for analyzing summary statistics, ACAT does not need the LD information that is often estimated from a population reference panel, which greatly speeds up the computation and avoids the potential issues caused by the estimation accuracy of the LD structure^29^. For example, it is very convenient and simple to use ACAT to perform gene-based analysis to complement the standard single-variant analysis in GWAS. The p-values from single-variant analysis can be directly used and are the only input required by ACAT for gene-based or pathway/network-based analysis, and therefore the computation can be done very efficiently.

As an omnibus testing procedure, ACAT in principle can be applied to combine complementary methods in nearly all kinds of genetic studies, including single-variant analysis, multiple-traits analysis and set-based analysis considered in this paper. In these studies, there often exists multiple competitive methods developed based on different reasonable assumptions. For instance, in set-based analysis, the assumptions of SKAT, ACAT-V and the burden test differ in the number and directionality of the casual variants. In multiple-trait analysis, the performance of different methods depends on many factors such as the number of traits associated with a variant and the heterogeneity of effect sizes. Due to the absence of prior knowledge about the underlying genetic architecture, omnibus testing can lead to robust analysis result and enhance the overall power. The ability of ACAT to obtain a p-value efficiently without simulation-based approaches allows for rapid combination of multiple methods and makes omnibus testing feasible for large studies even at the whole-genome scale.

While equal weights are employed in ACAT-O to combine different set-based tests, one can also consider upweighting the tests that are more likely to be powerful in a particular analysis to further boost the power. For example, if there are previous studies showing the existence of both protective and harmful variants for a trait, one can give less weight to the burden test and more weight to SKAT and ACAT-V. Hence, as the understanding of a trait progresses, the omnibus test constructed by ACAT has the capacity to mature to increase power. In contrast, the minimum p-value method and the Fisher’s method do not allow for flexible weights for the combination of tests.

The whole genome sequencing analysis of the ARIC data clearly demonstrates that the choices of weights can have a substantial impact on the power of a test. In the analysis results, SKAT with the default beta(MAF; 1,25) weights performs better for Lp(a) but identifies markedly fewer significant windows for neutrophil count than SKAT with the equal weights (or the beta(MAF; 1,1) weights). This also indicates that a single type of weights does not uniformly give the best performance across different studies and it is necessary to determine the weights in an adaptive manner based on the observed data. Besides MAF, one can also consider incorporating various functional annotations as weights for SKAT, ACAT-V and the burden test to further boost power, if the functional annotations are expected to be predictive for effect sizes and/or the probabilities of variants being causal. Since it is rarely known in advance that which functional annotation would lead to the optimal power, one can also use ACAT to combine the p-values of set-based tests weighted by multiple functional annotations that are potentially informative.

Another interesting observation from our analysis of the ARIC sequencing data is that even though some underpowered tests are included for omnibus testing, ACAT-O would only have little power loss. For example, the significant windows detected by the burden tests are only a small proportion of those detected by SKAT and ACAT-V in all the analyses. Including the burden tests in ACAT-O, however, resulted in little power loss. Hence, applying ACAT to combine multiple functional annotations could still be beneficial even if some non-informative functional annotations are incorporated.

## Supporting information

## Supplemental Data

Supplemental Data include nine figures and four tables.

## Acknowledgments

This work was supported by grants R35 CA197449, P01-CA134294, U01-HG009088, U19-CA203654, and R01-HL113338 (to X.L.). The Atherosclerosis Risk in Communities (ARIC) study is carried out as a collaborative study supported by the National Heart, Lung, and Blood Institute (NHLBI) contracts (HHSN268201100005C, HHSN268201100006C, HHSN268201100007C, HHSN268201100008C, HHSN268201100009C, HHSN268201100010C, HHSN268201100011C, and HHSN268201100012C). The authors thank the staff and participants of the ARIC study for their important contributions. Sequencing was carried out at the Baylor College of Medicine Human Genome Sequencing Center, also supported by the National Human Genome Research Institute grants U54 HG003273 and UM1 HG008898.

## Web Resources

The URLs for software programs referenced in this article are as follows: ACAT, *https://github.com/yaowuliu/ACAT*

## Appendix A. The approximation accuracy of the ACAT p-value

### The theory

The theory of ACAT was studied in details in Liu and Xie^19^, and we provide a brief description related to rare-variant analysis here. Recall that the ACAT test statistic is defined as 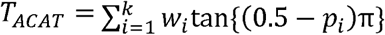, where p_i_’s are the p-values, w’s are nonnegative weights. We can transform the p-values to z-scores *z*_*i*_’s, i.e., *p*_*i*_ = 2{1-Φ(| z _i_|)} for i = 1,2,…,k. If each pair of the z-scores follows a bivariate normal distribution with mean 0 under the null hypothesis, then we have

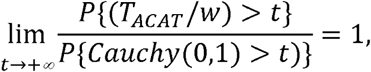

where 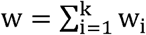 and *Cauchy*(0,1) denotes a random variable following the standard Cauchy distribution. This result holds under an arbitrary correlation structure of the z-scores/p-values and indicates that the tail of the ACAT test statistic is approximately Cauchy distributed. As p-value corresponds to the tail probability of the null distribution, the theoretical result suggests that the Cauchy distribution can be used to approximate the p-value of ACAT and the approximation would be particularly accurate when the ACAT p-value is very small. Because of the necessity of adopting stringent p-value threshold in sequencing studies to control for the rate of false-positive findings, the Cauchy-distribution-based approximation would be very accurate for the regions that are significantly associated with the trait. In some cases, the p-values combined by ACAT might not exactly follow a uniform distribution between 0 and 1 under the null hypothesis. For instance, the calibrated SKAT or variant-level p-value based on the analytic approximations is often conservative for very rare variants and dichotomous traits. When the p-values combined by ACAT are conservative, we have

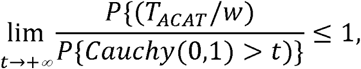

which indicates that the p-value of ACAT based on the Cauchy approximation would be also conservative.

### Practical guidelines

As ACAT is a general method for combing p-values and can be used for many other applications beyond variant set analysis in WGS, we provide guidelines regarding the accuracy of the ACAT p-value calculated by the Cauchy-distribution-based approximation. The guidelines are based on the assumption that the p-values aggregated by ACAT are accurate in the sense that they follow a uniform distribution between 0 and 1 under the null hypothesis. If this assumption is violated and the p-values are conservative, then the ACAT p-value generally would also be conservative as mentioned earlier.

The approximation accuracy certainly would depend on many factors, among which the ACAT p-value (i.e., *p*_ACAT_) itself and the correlation among the p-values are most important. In general, as implied by the theory, the smaller *p*_ACAT_ is, the less impact the correlation could have on the accuracy of *p*_ACAT_. When the ACAT p-value is very small (e.g., *p*_ACAT_ *<* 10^−5^), the type I error would be well controlled under almost all kinds of correlation structures, unless in a very rare situation that we will describe later. When the ACAT p-value is moderately small (e.g., 10^−3^ *< p*_ACAT_ *<* 10^−5^), the accuracy is generally satisfactory for practical use but a slight inflation is possible. When the ACAT p-value is large (e.g., *p_ACAT_* *<* 10^−3^), one may need to pay attention to the potential type I error inflation when the correlations are moderately strong.

A rare situation that one should be always cautious of is when there exist many strong negative correlations among the p-values. Fortunately, this situation seldomly happens in practice. For example, if the p-values are calculated from two-sided z-scores, it is impossible to have strong negative correlation between two p-values. Moreover, the p-values of competitive methods (e.g., the burden test, SKAT and ACAT-V) are also often positively correlated.

