## Supplementary material for "ACAT: A Fast and Powerful P-value Combination Method for Rare-variant Analysis in Sequencing Studies"

**Figure S1**

The genomic landscape of sliding windows significantly associated with Lp(a) levels among AAs on chromosome 6. Seven methods are compared: ACAT-V(1,1), ACAT-V(1,25), SKAT(1,1), SKAT(1,25), Burden(1,1), Burden(1,25) and the omnibus test ACAT-O that combines the other six tests, where the two numbers in the parentheses indicate the choice of the beta(MAF) weight parameters $a_1$ and $a_2$ in the test. A dot means that the sliding window at this location is significant by the method that the color of the dot represents. The numbers on the left of the plot show the number of significant windows identified by each method.

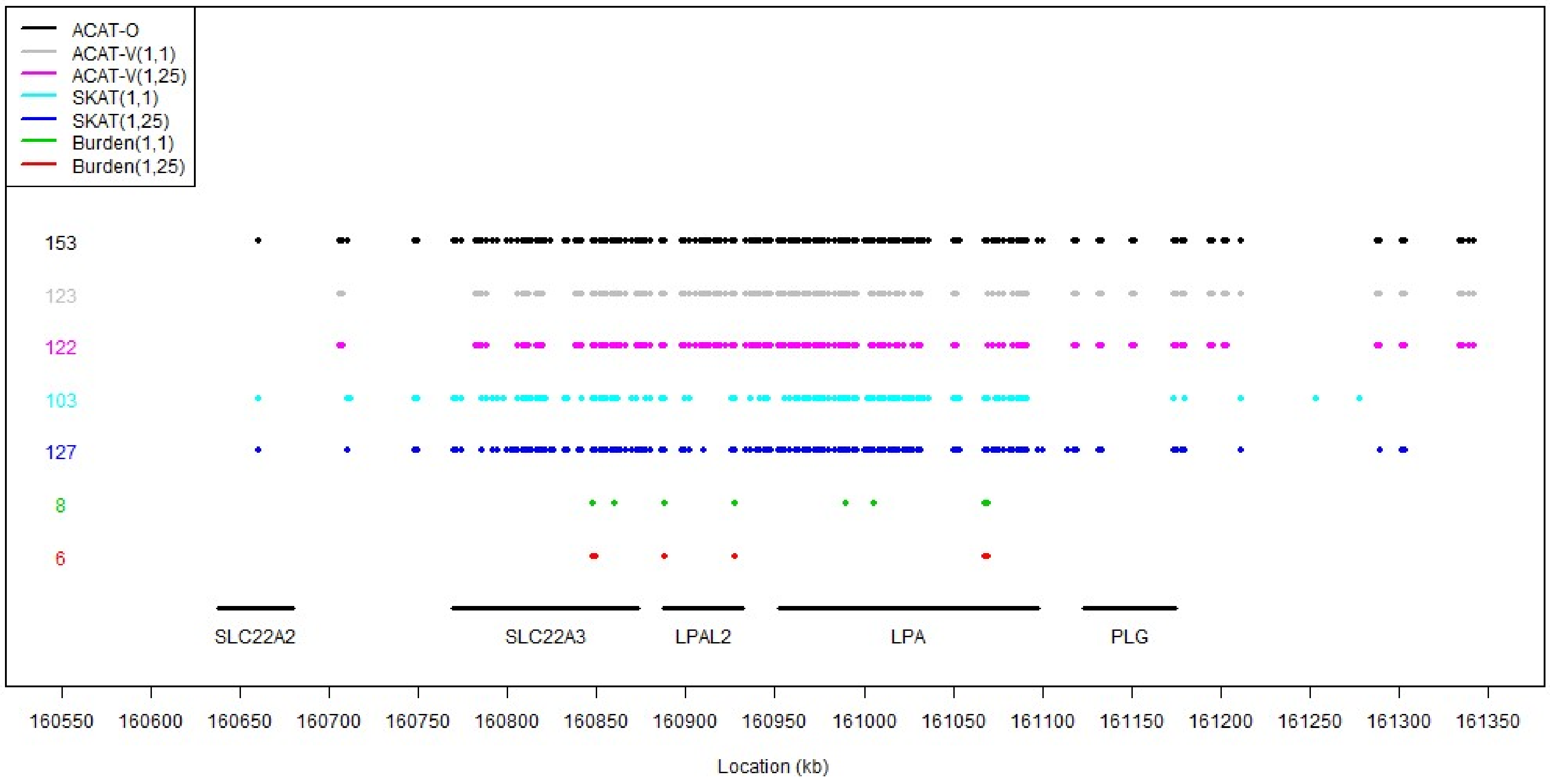

**Figure S2**

The genomic landscape of sliding windows significantly associated with Lp(a) levels among EAs on chromosome 6. Seven methods are compared: ACAT-V(1,1), ACAT-V(1,25), SKAT(1,1), SKAT(1,25), Burden(1,1), Burden(1,25) and the omnibus test ACAT-O that combines the other six tests, where the two numbers in the parentheses indicate the choice of the beta(MAF) weight parameters $a_1$ and $a_2$ in the test. A dot means that the sliding window at this location is significant by the method that the color of the dot represents. The numbers on the left of the plot show the number of significant windows identified by each method.

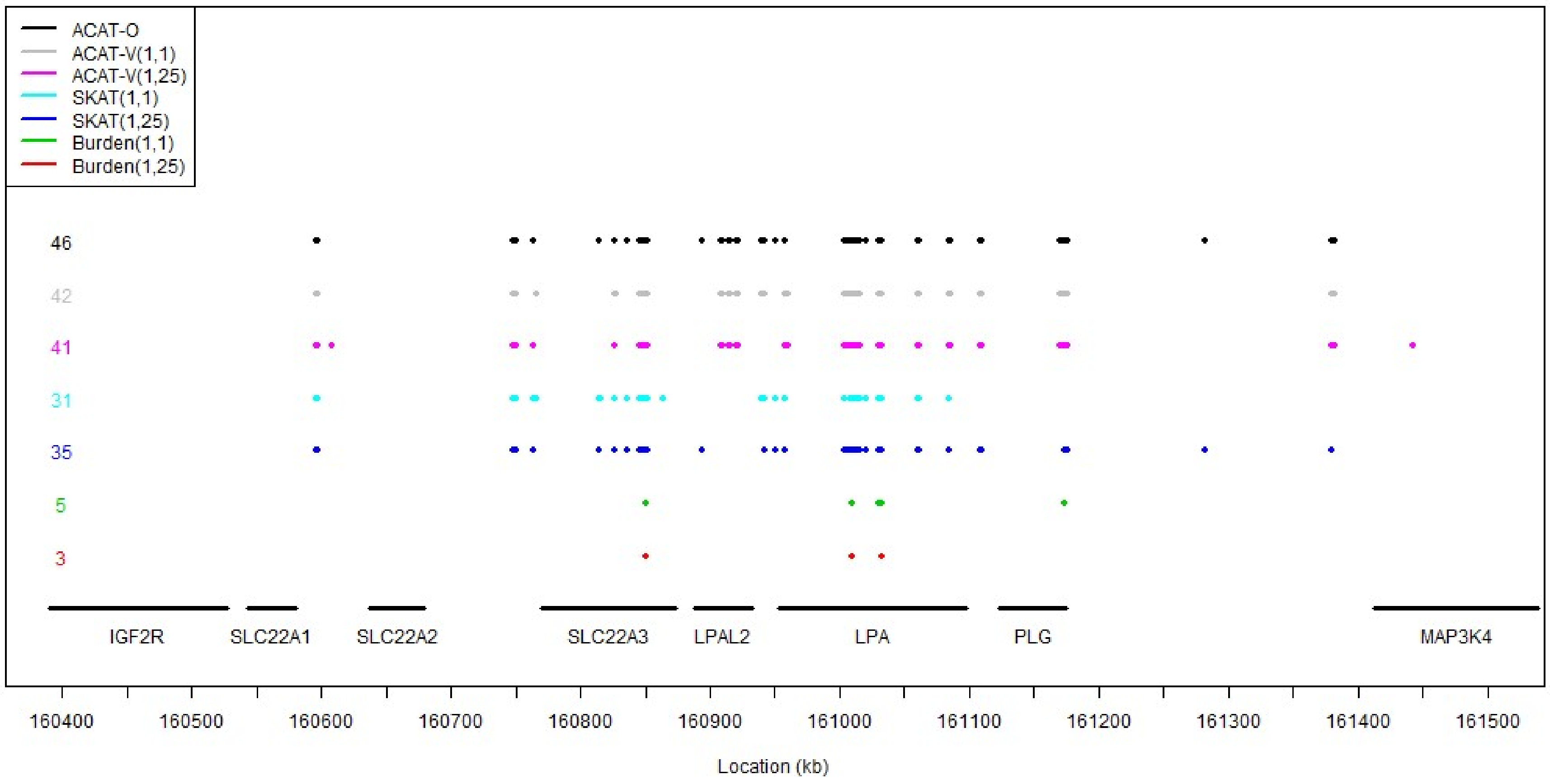

**Figure S3**

The genomic landscape of sliding windows significantly associated with neutrophil count among AAs on chromosome 1. Seven methods are compared: ACAT-V(1,1), ACAT-V(1,25), SKAT(1,1), SKAT(1,25), Burden(1,1), Burden(1,25) and the omnibus test ACAT-O that combines the other six tests, where the two numbers in the parentheses indicate the choice of the beta(MAF) weight parameters $a_1$ and $a_2$ in the test. A dot means that the sliding window at this location is significant by the method that the color of the dot represents. The numbers on the left of the plot show the number of significant windows identified by each method.

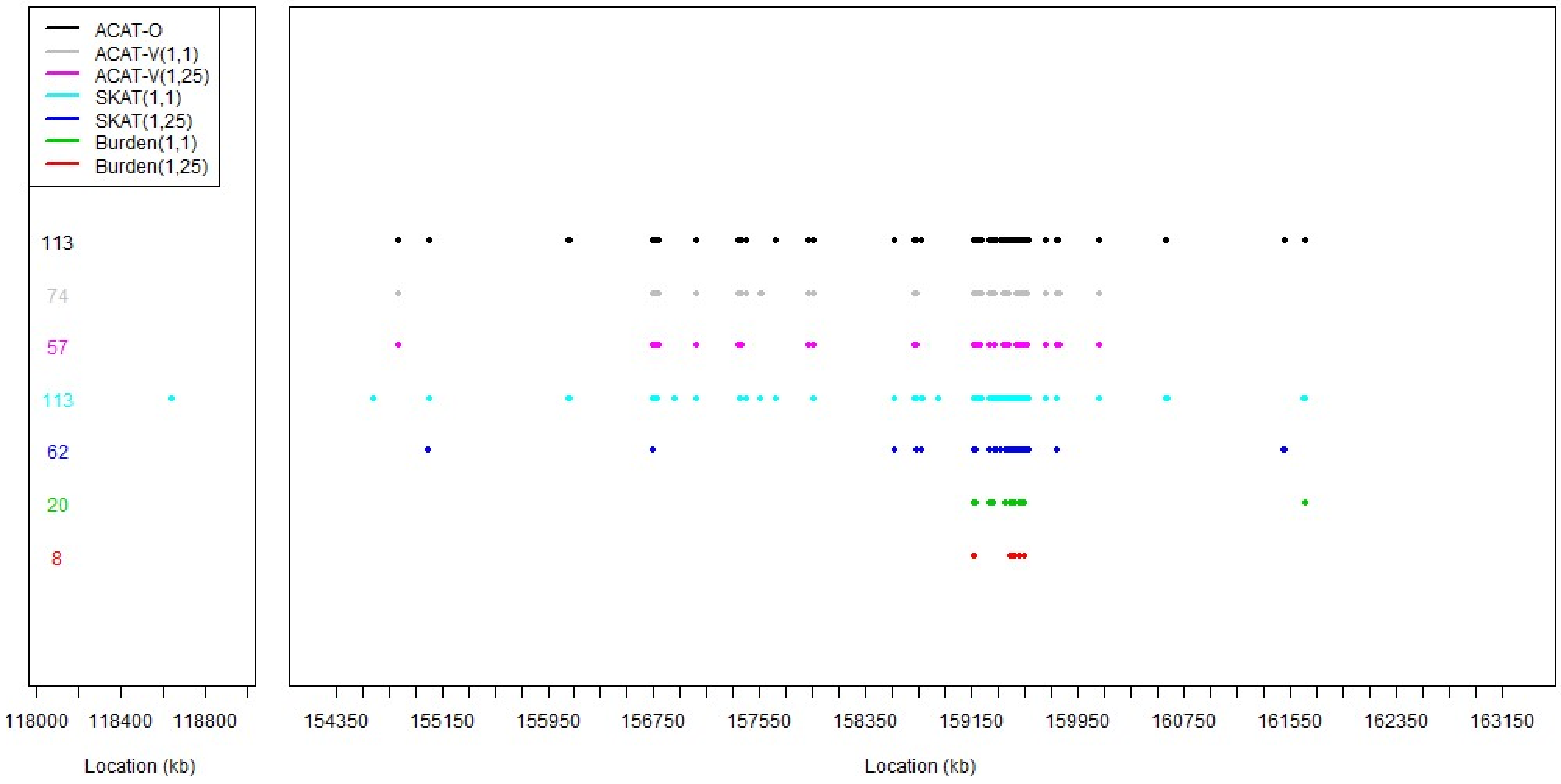

**Figure S4**

Scatterplot of p-values of significant sliding windows that are associated with Lp(a) levels among AAs (i.e., those reported in Table S2). The p-values are shown at the -log10 scale. The dashed lines in the plots correspond to the significance threshold $\alpha = 3.75 \times 10^{-8}$.

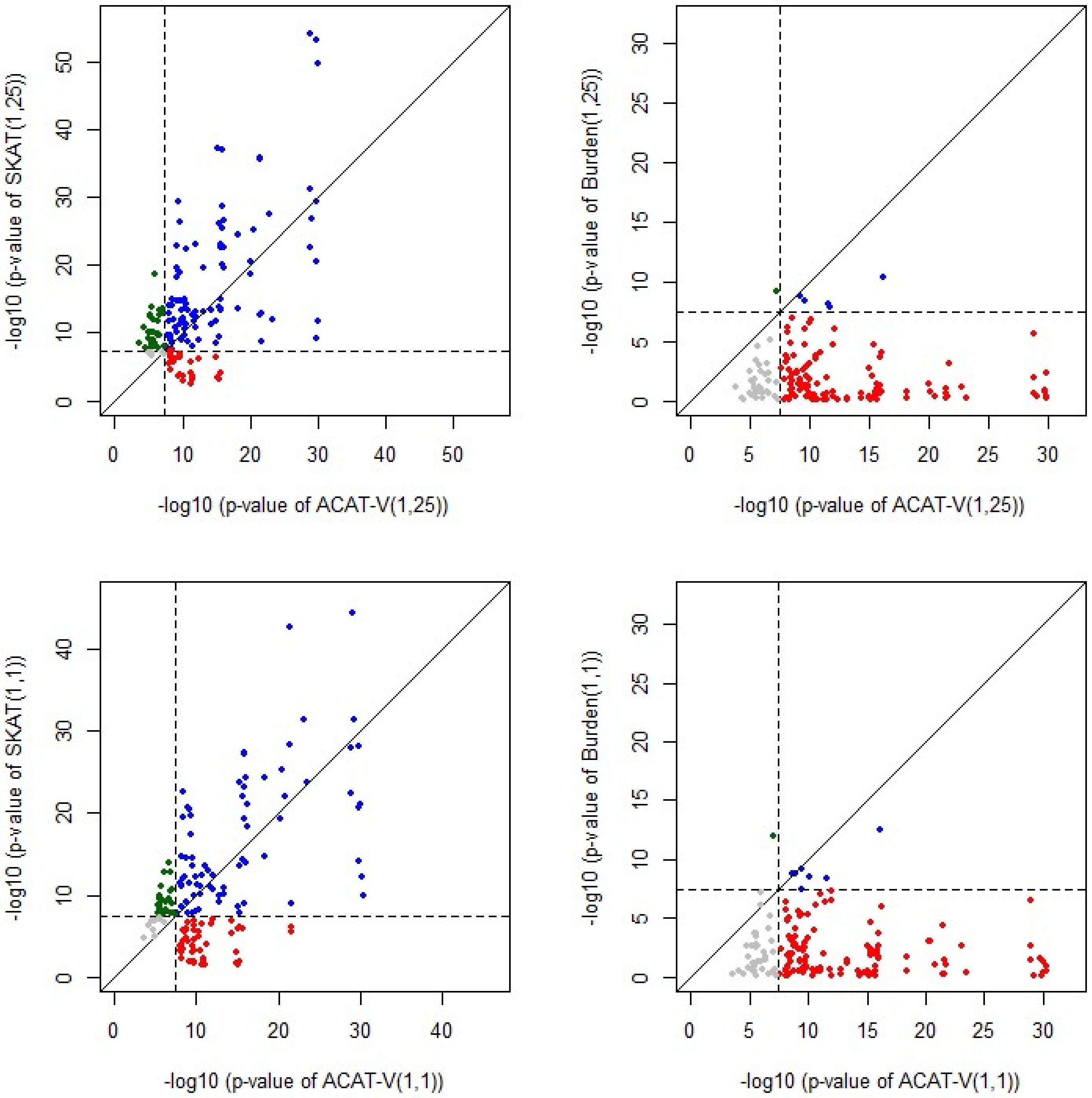

**Figure S5**

Scatterplot of p-values of significant sliding windows that are associated with Lp(a) levels among EAs (i.e., those reported in Table S3). The p-values are shown at the -log10 scale. The dashed lines in the plots correspond to the significance threshold $\alpha = 3.75 \times 10^{-8}$.

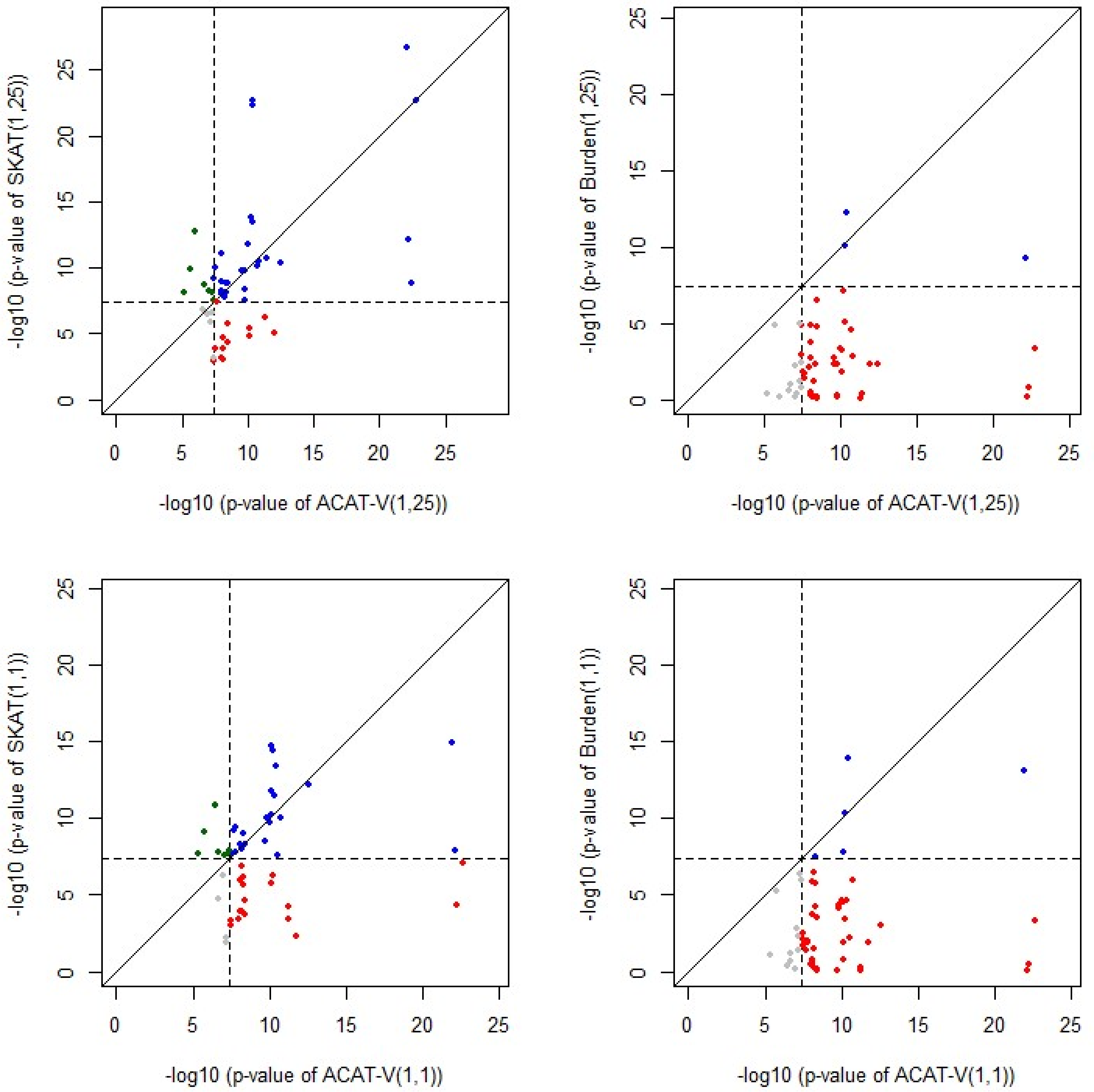

**Figure S6**

Scatterplot of p-values of significant sliding windows that are associated with neutrophil count among AAs (i.e., those reported in Table S4). The p-values are shown at the -log10 scale. The dashed lines in the plots correspond to the significance threshold $\alpha = 3.75 \times 10^{-8}$.

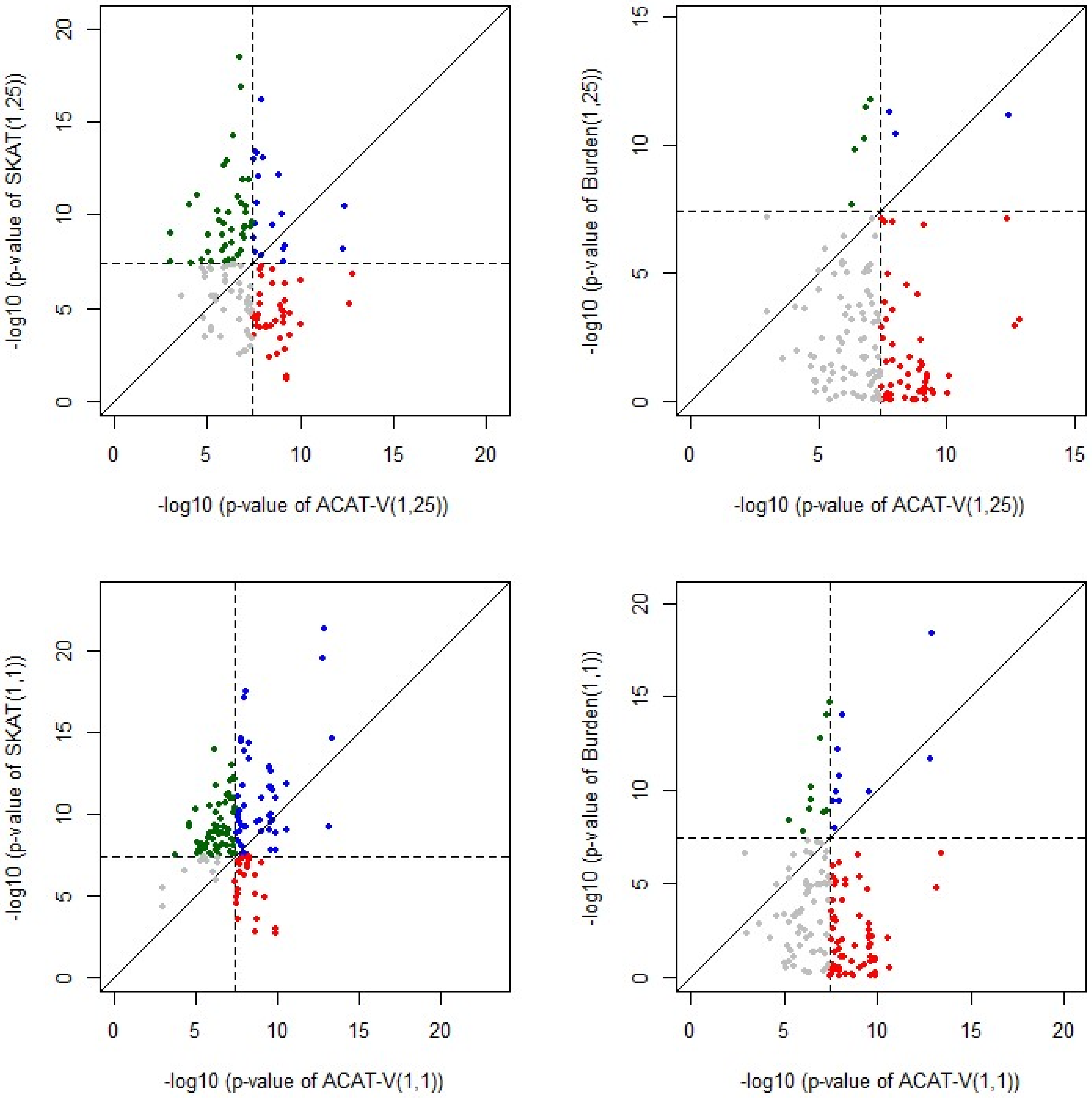

**Figure S7**

CADD-Phred score for the variants in sliding windows that are only identified by ACAT-V (not by SKAT or the burden test) as significantly associated with Lp(a) levels among AAs.

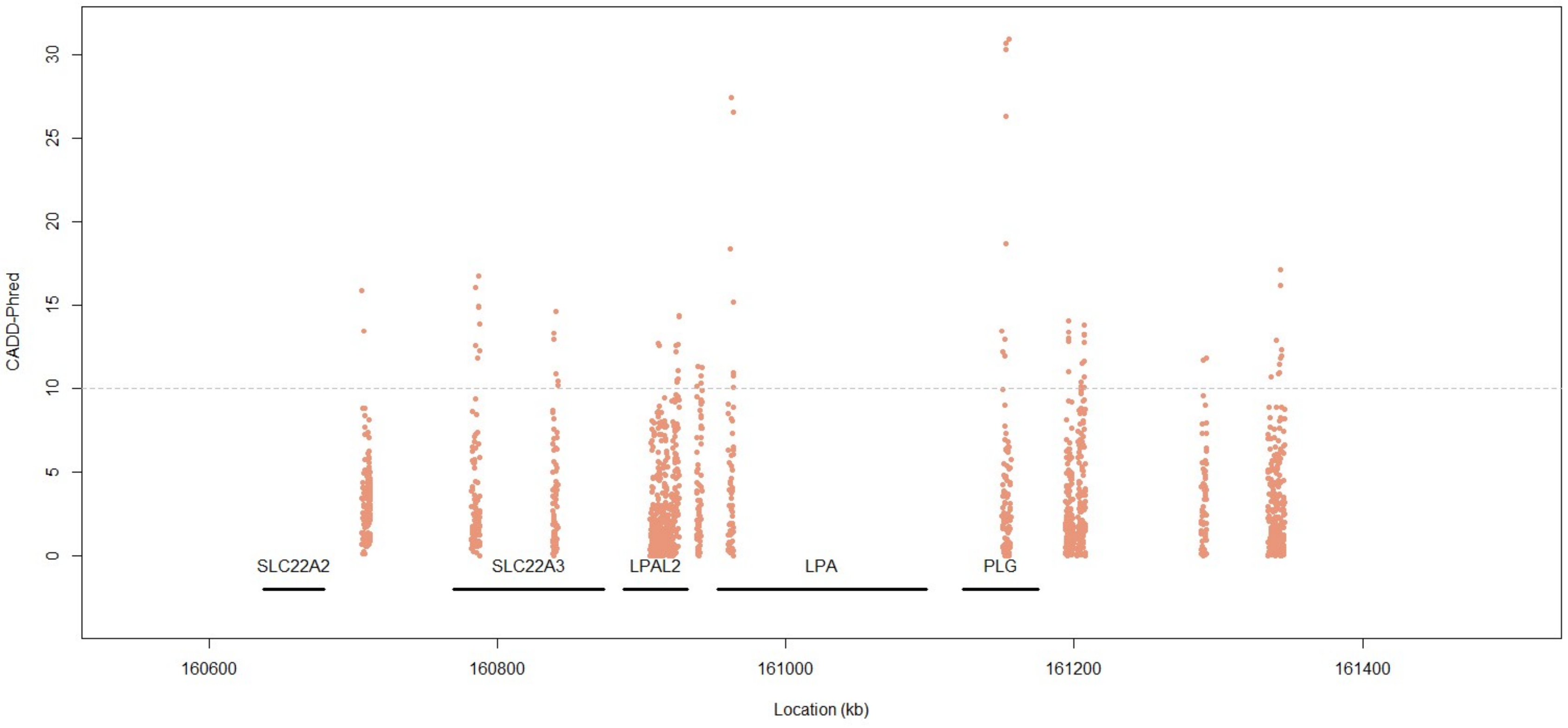

**Figure S8**

CADD-Phred score for the variants in sliding windows that are only identified by ACAT-V (not by SKAT or the burden test) as significantly associated with Lp(a) levels among EAs.

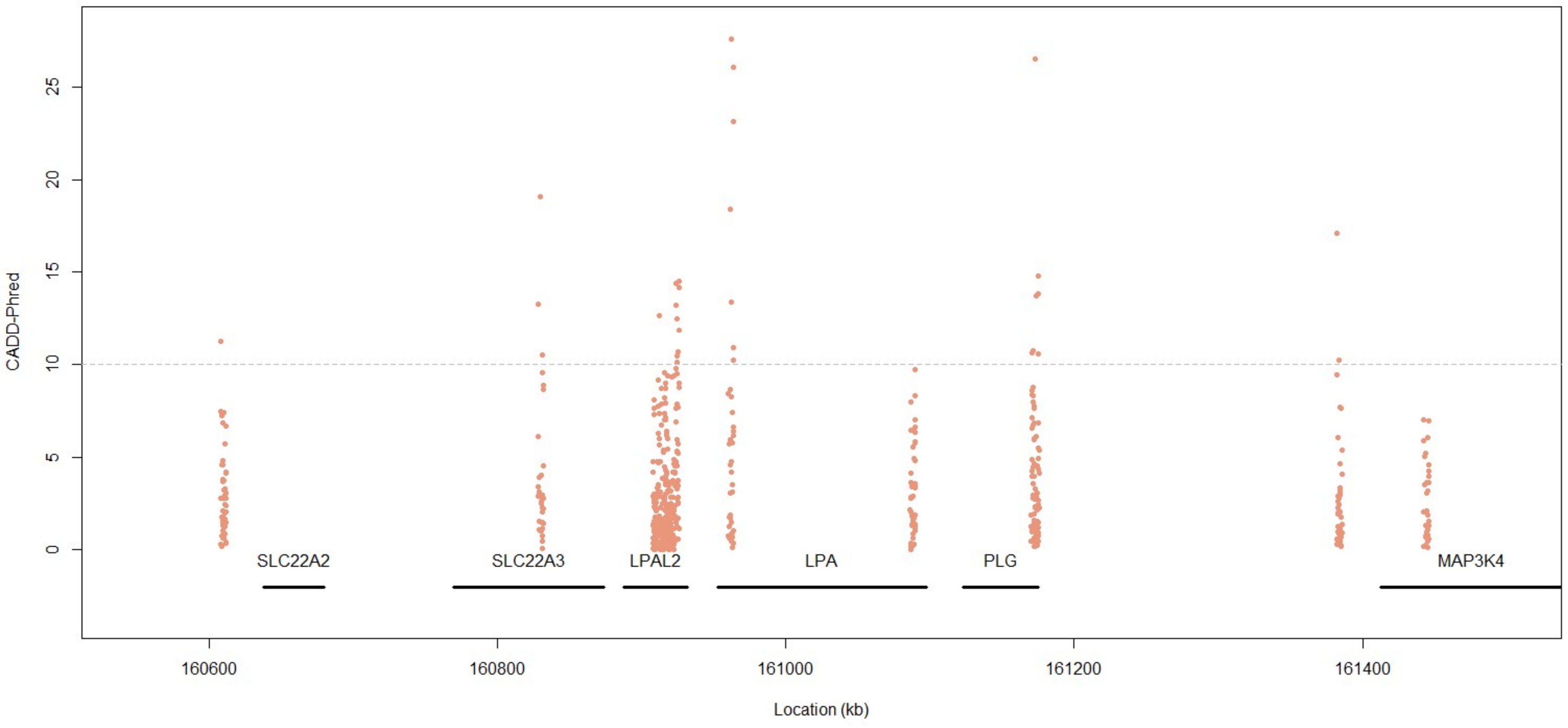

**Figure S9**

CADD-Phred score for the variants in sliding windows that are only identified by ACAT-V (not by SKAT or the burden test) as significantly associated with neutrophil count among AAs.

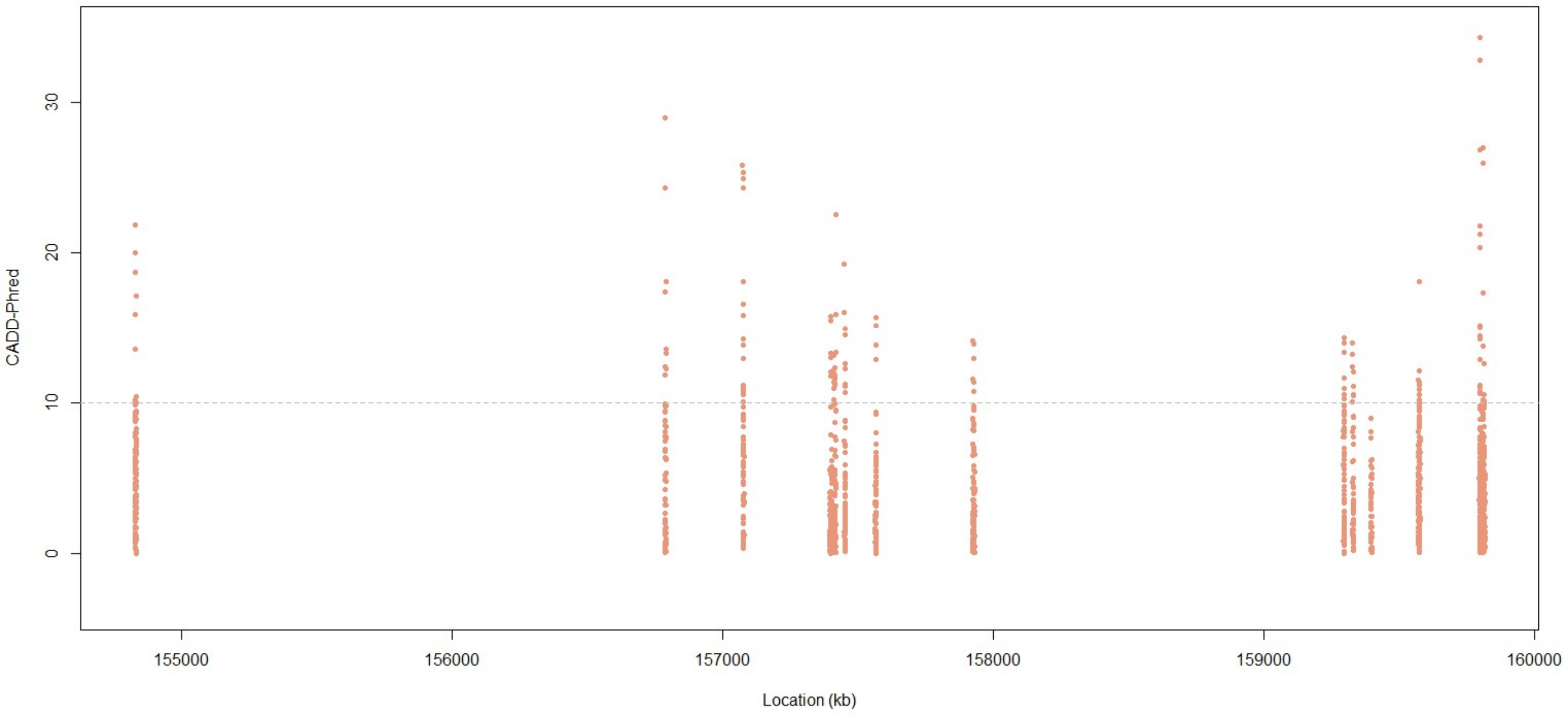

**Table S1**

The values of parameters $b$ and $c$ chosen in the power simulations, which depend on the percentage of casual variants in a SNV set.

| Percentage of casual variants | Continuous traits | | Dichotomous traits | |
|---|---|---|---|---|
| | $b$ | $c$ | $b$ | $c$ |
| 5% | 1.0 | 0.5 | 1.5 | 1.0 |
| 20% | 0.5 | 0.2 | 0.7 | 0.4 |
| 50% | 0.2 | 0.1 | 0.3 | 0.2 |

**Table S2**

Significant 4kb sliding windows for Lp(a) levels in AAs. The p-value threshold for genome-wide significance is $3.75 \times 10^{-8}$. Seven methods are compared: ACAT-V(1,1), ACAT-V(1,25), SKAT(1,1), SKAT(1,25), Burden(1,1), Burden(1,25) and the omnibus test ACAT-O that combines the other six tests, where the two numbers in the parentheses indicate the choice of the beta(MAF) weight parameters $a_1$ and $a_2$ in the test.

| Chr. | Start position (bp) | End position (bp) | cMAF | #SNV | P-values | | | | | | |
|---|---|---|---|---|---|---|---|---|---|---|---|
| | | | | | Burden (1,25) | Burden (1,1) | SKAT (1,25) | SKAT (1,1) | ACAT-V (1,25) | ACAT-V (1,1) | ACAT-O |
| 6 | 160660009 | 160664008 | 0.40 | 74 | 8.70E-02 | 8.07E-03 | 3.33E-09 | 7.49E-10 | 1.39E-06 | 6.52E-07 | 3.67E-09 |
| 6 | 160706009 | 160710008 | 0.40 | 56 | 2.77E-03 | 6.22E-04 | 3.27E-07 | 3.56E-06 | 4.82E-09 | 2.71E-09 | 1.04E-08 |
| 6 | 160708009 | 160712008 | 0.58 | 80 | 3.14E-04 | 3.48E-04 | 1.78E-07 | 2.30E-06 | 7.14E-09 | 4.02E-09 | 1.52E-08 |
| 6 | 160710009 | 160714008 | 0.74 | 87 | 2.34E-01 | 7.82E-02 | 7.48E-11 | 1.60E-09 | 9.73E-06 | 5.57E-06 | 4.29E-10 |
| 6 | 160712009 | 160716008 | 0.66 | 81 | 4.00E-03 | 2.53E-03 | 1.02E-07 | 1.95E-08 | 9.87E-06 | 5.96E-06 | 9.77E-08 |
| 6 | 160748009 | 160752008 | 0.53 | 61 | 6.64E-02 | 3.53E-04 | 1.28E-09 | 5.61E-09 | 4.78E-06 | 2.93E-06 | 6.26E-09 |
| 6 | 160750009 | 160754008 | 0.41 | 62 | 1.80E-02 | 1.94E-04 | 2.00E-14 | 2.00E-09 | 4.06E-06 | 2.60E-06 | 1.20E-13 |
| 6 | 160770009 | 160774008 | 0.33 | 58 | 6.20E-01 | 3.51E-02 | 3.52E-09 | 6.30E-09 | 9.96E-07 | 4.85E-07 | 1.35E-08 |
| 6 | 160772009 | 160776008 | 0.36 | 63 | 2.65E-01 | 5.63E-01 | 2.24E-13 | 3.05E-10 | 2.72E-07 | 2.18E-07 | 1.35E-12 |
| 6 | 160774009 | 160778008 | 0.44 | 82 | 6.31E-03 | 2.11E-02 | 1.42E-08 | 8.54E-09 | 4.96E-07 | 3.88E-07 | 3.12E-08 |
| 6 | 160782009 | 160786008 | 0.26 | 52 | 6.35E-01 | 3.34E-01 | 3.05E-06 | 5.33E-04 | 8.54E-09 | 8.70E-09 | 2.58E-08 |
| 6 | 160784009 | 160788008 | 0.36 | 53 | 6.03E-01 | 8.64E-01 | 6.01E-08 | 1.60E-04 | 4.85E-09 | 6.33E-09 | 1.58E-08 |
| 6 | 160786009 | 160790008 | 0.42 | 65 | 9.47E-01 | 6.84E-01 | 1.49E-09 | 8.67E-10 | 4.47E-13 | 1.65E-13 | 7.22E-13 |
| 6 | 160788009 | 160792008 | 0.39 | 61 | 7.91E-01 | 2.73E-01 | 5.51E-07 | 1.01E-10 | 4.68E-13 | 1.53E-13 | 6.91E-13 |
| 6 | 160792009 | 160796008 | 0.37 | 63 | 5.41E-03 | 1.01E-03 | 2.68E-10 | 1.88E-13 | 3.23E-07 | 1.20E-07 | 1.13E-12 |
| 6 | 160794009 | 160798008 | 0.32 | 49 | 2.25E-02 | 8.98E-03 | 2.54E-10 | 1.15E-09 | 2.68E-07 | 1.07E-07 | 1.24E-09 |
| 6 | 160798009 | 160802008 | 0.28 | 59 | 6.20E-01 | 3.02E-01 | 1.84E-07 | 1.32E-08 | 3.40E-06 | 5.59E-06 | 7.33E-08 |
| 6 | 160800009 | 160804008 | 0.36 | 58 | 1.10E-03 | 1.56E-03 | 6.42E-11 | 5.14E-08 | 1.89E-06 | 2.79E-06 | 3.85E-10 |
| 6 | 160802009 | 160806008 | 0.37 | 65 | 2.69E-05 | 2.32E-04 | 1.01E-09 | 8.18E-08 | 2.50E-06 | 3.31E-06 | 5.96E-09 |
| 6 | 160804009 | 160808008 | 0.42 | 74 | 3.70E-01 | 2.28E-02 | 3.23E-08 | 1.74E-07 | 8.16E-06 | 1.47E-05 | 1.63E-07 |
| 6 | 160806009 | 160810008 | 0.38 | 77 | 7.17E-01 | 2.75E-01 | 2.38E-10 | 5.07E-12 | 1.34E-08 | 6.94E-09 | 2.97E-11 |
| 6 | 160808009 | 160812008 | 0.28 | 66 | 1.54E-04 | 9.07E-06 | 1.83E-12 | 8.81E-13 | 9.65E-09 | 4.77E-09 | 3.57E-12 |
| 6 | 160810009 | 160814008 | 0.32 | 55 | 6.19E-04 | 1.17E-02 | 1.11E-14 | 2.95E-20 | 1.03E-08 | 4.08E-09 | 1.11E-16 |
| 6 | 160812009 | 160816008 | 0.39 | 51 | 1.61E-02 | 1.57E-01 | 1.47E-13 | 2.80E-23 | 1.20E-08 | 5.20E-09 | 1.11E-16 |
| 6 | 160814009 | 160818008 | 0.41 | 56 | 1.89E-01 | 6.03E-02 | 1.47E-12 | 3.09E-08 | 1.67E-06 | 1.22E-06 | 8.83E-12 |
| 6 | 160816009 | 160820008 | 0.51 | 68 | 6.97E-07 | 4.74E-07 | 1.86E-10 | 3.65E-12 | 6.80E-09 | 8.61E-09 | 2.15E-11 |
| 6 | 160818009 | 160822008 | 0.53 | 68 | 1.58E-06 | 2.08E-06 | 9.12E-10 | 2.89E-09 | 6.55E-09 | 6.95E-09 | 3.45E-09 |
| 6 | 160820009 | 160824008 | 0.37 | 58 | 1.87E-03 | 5.08E-03 | 7.71E-09 | 1.54E-08 | 2.51E-08 | 2.08E-08 | 2.12E-08 |
| 6 | 160822009 | 160826008 | 0.38 | 54 | 7.20E-02 | 8.27E-01 | 1.02E-08 | 5.69E-09 | 1.69E-06 | 3.15E-06 | 2.19E-08 |
| 6 | 160824009 | 160828008 | 0.39 | 48 | 2.15E-02 | 2.16E-03 | 5.91E-10 | 2.01E-06 | 7.73E-06 | 1.57E-05 | 3.55E-09 |
| 6 | 160826009 | 160830008 | 0.30 | 38 | 3.10E-01 | 1.64E-01 | 1.48E-08 | 1.23E-05 | 6.33E-06 | 1.26E-05 | 8.81E-08 |
| 6 | 160832009 | 160836008 | 0.19 | 49 | 7.58E-06 | 7.04E-06 | 4.53E-14 | 3.62E-11 | 2.11E-07 | 1.50E-07 | 2.72E-13 |
| 6 | 160834009 | 160838008 | 0.25 | 48 | 1.99E-01 | 2.50E-02 | 1.88E-12 | 6.59E-09 | 4.18E-07 | 4.78E-07 | 1.13E-11 |
| 6 | 160838009 | 160842008 | 0.45 | 62 | 2.31E-02 | 1.83E-02 | 2.34E-06 | 5.20E-05 | 2.58E-09 | 3.68E-09 | 9.10E-09 |
| 6 | 160840009 | 160844008 | 0.41 | 60 | 4.70E-03 | 1.28E-03 | 1.40E-11 | 1.66E-06 | 1.35E-09 | 1.70E-09 | 8.24E-11 |
| 6 | 160842009 | 160846008 | 0.35 | 56 | 1.02E-01 | 4.81E-02 | 8.77E-15 | 6.04E-13 | 2.69E-09 | 2.81E-09 | 5.20E-14 |
| 6 | 160848009 | 160852008 | 0.17 | 44 | 1.52E-08 | 3.88E-09 | 1.84E-12 | 1.08E-11 | 2.11E-12 | 2.40E-12 | 3.94E-12 |
| 6 | 160850009 | 160854008 | 0.28 | 45 | 8.27E-09 | 4.14E-07 | 2.28E-13 | 1.15E-13 | 2.73E-12 | 4.22E-12 | 4.37E-13 |
| 6 | 160852009 | 160856008 | 0.35 | 60 | 6.82E-01 | 6.75E-02 | 3.17E-20 | 7.31E-11 | 6.66E-14 | 4.88E-14 | 1.11E-16 |
| 6 | 160854009 | 160858008 | 0.32 | 56 | 7.80E-01 | 4.36E-02 | 8.50E-14 | 1.36E-11 | 6.42E-14 | 4.35E-14 | 1.19E-13 |
| 6 | 160856009 | 160860008 | 0.28 | 53 | 4.09E-02 | 5.75E-02 | 6.36E-20 | 4.95E-18 | 4.38E-10 | 5.09E-10 | 3.77E-19 |
| 6 | 160858009 | 160862008 | 0.31 | 54 | 1.69E-03 | 5.25E-03 | 1.89E-15 | 2.55E-15 | 2.32E-10 | 2.54E-10 | 6.66E-15 |
| 6 | 160860009 | 160864008 | 0.28 | 54 | 1.44E-07 | 2.82E-09 | 1.13E-12 | 4.85E-12 | 8.27E-11 | 8.12E-11 | 5.38E-12 |
| 6 | 160862009 | 160866008 | 0.33 | 49 | 7.53E-04 | 5.02E-04 | 7.68E-13 | 7.34E-13 | 9.74E-11 | 1.20E-10 | 2.24E-12 |
| 6 | 160864009 | 160868008 | 0.31 | 56 | 8.08E-03 | 5.23E-02 | 1.09E-09 | 6.51E-07 | 1.73E-10 | 2.28E-10 | 5.41E-10 |
| 6 | 160866009 | 160870008 | 0.22 | 50 | 6.82E-02 | 1.24E-01 | 1.14E-12 | 2.30E-06 | 1.60E-10 | 1.57E-10 | 6.76E-12 |
| 6 | 160870009 | 160874008 | 0.25 | 52 | 3.77E-01 | 4.98E-02 | 3.09E-09 | 7.90E-09 | 4.13E-06 | 1.83E-06 | 1.33E-08 |
| 6 | 160872009 | 160876008 | 0.48 | 65 | 5.58E-02 | 3.73E-03 | 5.10E-14 | 1.09E-10 | 1.39E-10 | 1.83E-10 | 3.06E-13 |
| 6 | 160874009 | 160878008 | 0.44 | 70 | 4.27E-01 | 2.04E-01 | 1.73E-12 | 1.77E-07 | 1.28E-10 | 1.61E-10 | 1.02E-11 |
| 6 | 160876009 | 160880008 | 0.31 | 64 | 9.25E-01 | 8.20E-01 | 2.09E-10 | 3.10E-06 | 4.22E-11 | 4.48E-11 | 1.18E-10 |
| 6 | 160878009 | 160882008 | 0.24 | 51 | 3.42E-04 | 1.13E-04 | 6.44E-15 | 6.52E-11 | 3.21E-11 | 3.07E-11 | 3.86E-14 |
| 6 | 160880009 | 160884008 | 0.14 | 39 | 3.01E-07 | 5.38E-06 | 1.89E-15 | 8.96E-09 | 1.05E-10 | 1.15E-10 | 1.15E-14 |
| 6 | 160886009 | 160890008 | 0.56 | 78 | 1.74E-05 | 5.24E-06 | 6.72E-12 | 1.88E-08 | 2.75E-10 | 3.33E-10 | 3.86E-11 |
| 6 | 160888009 | 160892008 | 0.53 | 69 | 4.56E-09 | 7.80E-10 | 1.02E-19 | 3.35E-14 | 2.59E-10 | 3.20E-10 | 1.11E-16 |
| 6 | 160898009 | 160902008 | 0.54 | 83 | 1.20E-02 | 1.08E-02 | 1.25E-08 | 1.20E-04 | 4.20E-12 | 4.90E-12 | 1.36E-11 |
| 6 | 160900009 | 160904008 | 0.42 | 75 | 1.08E-06 | 3.22E-07 | 1.12E-23 | 2.34E-11 | 8.91E-13 | 9.31E-13 | 1.11E-16 |
| 6 | 160902009 | 160906008 | 0.35 | 81 | 1.77E-05 | 5.33E-08 | 8.62E-14 | 3.92E-13 | 1.10E-12 | 1.02E-12 | 3.74E-13 |
| 6 | 160906009 | 160910008 | 2.00 | 104 | 3.12E-01 | 2.66E-01 | 2.98E-03 | 3.38E-02 | 6.39E-12 | 1.88E-11 | 2.86E-11 |
| 6 | 160908009 | 160912008 | 1.57 | 101 | 7.41E-01 | 3.79E-01 | 7.58E-05 | 1.11E-02 | 6.03E-12 | 1.47E-11 | 2.56E-11 |
| 6 | 160910009 | 160914008 | 0.96 | 85 | 3.90E-02 | 8.56E-01 | 1.66E-09 | 7.84E-04 | 2.04E-11 | 3.72E-11 | 7.84E-11 |
| 6 | 160912009 | 160916008 | 1.45 | 96 | 6.21E-01 | 3.20E-01 | 2.06E-06 | 6.11E-03 | 4.78E-12 | 1.10E-11 | 2.00E-11 |
| 6 | 160914009 | 160918008 | 1.98 | 107 | 2.48E-01 | 2.03E-01 | 2.75E-04 | 2.83E-02 | 3.72E-12 | 9.28E-12 | 1.59E-11 |
| 6 | 160916009 | 160920008 | 1.84 | 93 | 7.35E-01 | 3.52E-01 | 9.19E-04 | 3.96E-02 | 3.48E-12 | 8.61E-12 | 1.49E-11 |
| 6 | 160918009 | 160922008 | 1.43 | 77 | 4.75E-01 | 2.97E-01 | 7.32E-04 | 3.17E-02 | 3.33E-16 | 8.88E-16 | 1.55E-15 |

| | | | | | | | | | | | |
|---|---|---|---|---|---|---|---|---|---|---|---|
| 6 | 160920009 | 160924008 | 1.09 | 76 | 9.59E-01 | 5.65E-01 | 8.26E-05 | 1.21E-02 | 2.29E-16 | 4.44E-16 | 8.88E-16 |
| 6 | 160922009 | 160926008 | 1.04 | 75 | 7.93E-01 | 4.22E-01 | 3.82E-04 | 2.66E-02 | 7.77E-16 | 1.44E-15 | 3.11E-15 |
| 6 | 160926009 | 160930008 | 0.40 | 47 | 2.23E-04 | 2.58E-03 | 7.84E-21 | 1.23E-09 | 1.08E-16 | 1.16E-16 | 1.11E-16 |
| 6 | 160928009 | 160932008 | 0.24 | 40 | 4.70E-11 | 3.00E-13 | 2.13E-23 | 4.79E-19 | 6.45E-17 | 6.59E-17 | 1.28E-22 |
| 6 | 160934009 | 160938008 | 0.28 | 51 | 2.20E-03 | 2.59E-03 | 3.29E-09 | 1.43E-06 | 8.88E-16 | 8.88E-16 | 2.89E-15 |
| 6 | 160936009 | 160940008 | 0.38 | 68 | 2.05E-05 | 2.69E-05 | 1.63E-14 | 1.97E-08 | 4.44E-16 | 4.44E-16 | 1.33E-15 |
| 6 | 160938009 | 160942008 | 0.47 | 64 | 4.26E-01 | 4.57E-01 | 4.37E-07 | 1.18E-03 | 8.88E-16 | 1.11E-15 | 3.11E-15 |
| 6 | 160940009 | 160944008 | 0.37 | 52 | 3.78E-02 | 3.71E-01 | 5.38E-14 | 1.40E-06 | 2.19E-16 | 2.18E-16 | 6.66E-16 |
| 6 | 160942009 | 160946008 | 0.28 | 54 | 1.55E-01 | 6.00E-03 | 1.42E-37 | 4.81E-28 | 1.59E-16 | 1.23E-16 | 8.50E-37 |
| 6 | 160944009 | 160948008 | 0.21 | 42 | 8.61E-03 | 4.83E-03 | 6.72E-38 | 2.09E-24 | 5.55E-16 | 4.44E-16 | 4.03E-37 |
| 6 | 160946009 | 160950008 | 0.27 | 52 | 3.47E-01 | 1.51E-02 | 7.03E-27 | 3.11E-14 | 3.33E-16 | 4.44E-16 | 1.11E-16 |
| 6 | 160948009 | 160952008 | 0.30 | 59 | 1.88E-01 | 8.18E-03 | 4.36E-10 | 7.66E-07 | 3.33E-16 | 4.44E-16 | 1.22E-15 |
| 6 | 160952009 | 160956008 | 0.45 | 70 | 6.68E-01 | 7.58E-01 | 6.91E-12 | 5.57E-06 | 4.33E-15 | 4.44E-15 | 1.32E-14 |
| 6 | 160954009 | 160958008 | 0.52 | 78 | 2.79E-01 | 3.22E-01 | 3.72E-14 | 1.41E-07 | 4.44E-15 | 5.11E-15 | 1.34E-14 |
| 6 | 160956009 | 160960008 | 0.37 | 76 | 9.49E-02 | 1.21E-01 | 1.11E-15 | 5.88E-15 | 2.26E-16 | 2.05E-16 | 6.66E-16 |
| 6 | 160958009 | 160962008 | 0.37 | 60 | 8.24E-01 | 8.00E-01 | 2.52E-23 | 1.13E-22 | 2.29E-16 | 2.14E-16 | 1.23E-22 |
| 6 | 160960009 | 160964008 | 0.40 | 55 | 1.33E-02 | 9.97E-03 | 5.03E-08 | 1.95E-06 | 3.47E-09 | 2.06E-09 | 7.56E-09 |
| 6 | 160962009 | 160966008 | 0.32 | 52 | 1.63E-01 | 2.58E-02 | 2.50E-27 | 1.11E-14 | 9.56E-17 | 9.09E-17 | 1.11E-16 |
| 6 | 160964009 | 160968008 | 0.29 | 57 | 9.89E-05 | 1.11E-06 | 2.99E-20 | 8.87E-22 | 7.78E-17 | 5.36E-17 | 5.17E-21 |
| 6 | 160966009 | 160970008 | 0.34 | 74 | 3.70E-02 | 1.32E-02 | 2.62E-14 | 4.72E-20 | 1.99E-16 | 1.37E-16 | 1.11E-16 |
| 6 | 160968009 | 160972008 | 0.25 | 59 | 2.38E-01 | 1.94E-02 | 2.28E-29 | 5.45E-25 | 1.20E-16 | 1.03E-16 | 1.37E-28 |
| 6 | 160970009 | 160974008 | 0.28 | 48 | 5.01E-02 | 1.89E-04 | 4.64E-26 | 7.54E-24 | 1.25E-16 | 1.20E-16 | 2.76E-25 |
| 6 | 160972009 | 160976008 | 0.39 | 59 | 5.68E-01 | 9.24E-01 | 1.66E-12 | 2.85E-09 | 8.88E-16 | 6.66E-16 | 2.33E-15 |
| 6 | 160974009 | 160978008 | 0.37 | 61 | 1.13E-01 | 7.08E-03 | 1.03E-23 | 7.37E-28 | 1.72E-16 | 1.47E-16 | 4.42E-27 |
| 6 | 160976009 | 160980008 | 0.38 | 65 | 5.82E-01 | 6.66E-01 | 2.84E-36 | 2.13E-43 | 3.59E-22 | 3.09E-22 | 1.28E-42 |
| 6 | 160978009 | 160982008 | 0.52 | 73 | 9.45E-02 | 4.92E-05 | 1.63E-36 | 4.57E-29 | 3.64E-22 | 3.46E-22 | 9.78E-36 |
| 6 | 160980009 | 160984008 | 0.44 | 66 | 4.83E-01 | 1.11E-01 | 8.10E-26 | 1.13E-22 | 2.78E-21 | 1.51E-21 | 4.85E-25 |
| 6 | 160984009 | 160988008 | 0.63 | 84 | 1.54E-01 | 2.89E-02 | 3.53E-21 | 2.41E-21 | 1.82E-30 | 1.64E-30 | 5.18E-30 |
| 6 | 160986009 | 160990008 | 0.43 | 75 | 4.09E-01 | 4.23E-01 | 4.06E-30 | 6.67E-29 | 1.32E-30 | 1.12E-30 | 3.13E-30 |
| 6 | 160988009 | 160992008 | 0.36 | 78 | 1.51E-04 | 2.52E-03 | 9.47E-13 | 8.53E-10 | 2.39E-09 | 1.64E-09 | 5.67E-12 |
| 6 | 160990009 | 160994008 | 0.46 | 84 | 1.21E-07 | 1.76E-09 | 2.67E-09 | 2.55E-15 | 2.98E-09 | 2.17E-09 | 1.53E-14 |
| 6 | 160992009 | 160996008 | 0.54 | 72 | 1.32E-01 | 2.85E-01 | 1.05E-10 | 3.61E-12 | 5.86E-10 | 7.62E-10 | 2.07E-11 |
| 6 | 160994009 | 160998008 | 0.46 | 55 | 3.42E-02 | 9.49E-04 | 1.89E-19 | 4.11E-20 | 1.01E-20 | 5.01E-21 | 1.83E-20 |
| 6 | 160996009 | 161000008 | 0.33 | 51 | 2.02E-01 | 1.10E-03 | 3.54E-21 | 5.82E-26 | 7.95E-21 | 3.56E-21 | 3.49E-25 |
| 6 | 161000009 | 161004008 | 0.49 | 73 | 6.91E-02 | 6.68E-01 | 2.93E-09 | 1.68E-05 | 1.93E-04 | 2.84E-04 | 1.75E-08 |
| 6 | 161002009 | 161006008 | 0.55 | 73 | 2.73E-02 | 4.90E-01 | 3.43E-14 | 1.56E-08 | 6.61E-08 | 1.44E-07 | 2.06E-13 |
| 6 | 161004009 | 161008008 | 0.52 | 71 | 1.46E-05 | 1.41E-05 | 1.11E-15 | 2.44E-15 | 3.59E-09 | 6.38E-09 | 4.66E-15 |
| 6 | 161006009 | 161010008 | 0.58 | 79 | 1.00E-06 | 3.57E-08 | 4.36E-27 | 1.47E-08 | 3.00E-10 | 3.52E-10 | 1.11E-16 |
| 6 | 161008009 | 161012008 | 0.46 | 77 | 2.04E-01 | 8.24E-01 | 6.52E-54 | 6.22E-15 | 1.51E-30 | 1.39E-30 | 3.91E-53 |
| 6 | 161010009 | 161014008 | 0.34 | 70 | 4.29E-03 | 4.52E-02 | 1.90E-50 | 1.13E-21 | 1.11E-30 | 1.00E-30 | 1.14E-49 |
| 6 | 161012009 | 161016008 | 0.43 | 78 | 5.38E-01 | 3.27E-01 | 2.15E-14 | 2.33E-15 | 7.17E-19 | 3.79E-19 | 1.49E-18 |
| 6 | 161014009 | 161018008 | 0.45 | 71 | 1.57E-01 | 2.07E-02 | 2.95E-25 | 4.83E-25 | 6.63E-19 | 4.06E-19 | 1.10E-24 |
| 6 | 161016009 | 161020008 | 0.32 | 69 | 3.20E-02 | 2.13E-03 | 2.90E-13 | 8.02E-12 | 6.47E-06 | 2.33E-06 | 1.68E-12 |
| 6 | 161018009 | 161022008 | 0.34 | 76 | 2.71E-02 | 2.52E-06 | 3.05E-20 | 4.02E-21 | 6.73E-10 | 7.50E-10 | 2.13E-20 |
| 6 | 161020009 | 161024008 | 0.46 | 64 | 7.96E-02 | 6.66E-06 | 6.03E-30 | 2.39E-20 | 3.93E-10 | 5.30E-10 | 3.62E-29 |
| 6 | 161022009 | 161026008 | 0.57 | 62 | 4.02E-02 | 3.46E-04 | 1.49E-23 | 1.80E-21 | 8.69E-10 | 1.39E-09 | 8.84E-23 |
| 6 | 161024009 | 161028008 | 0.49 | 78 | 6.57E-04 | 7.24E-05 | 1.11E-10 | 1.23E-14 | 3.48E-07 | 2.02E-07 | 7.39E-14 |
| 6 | 161026009 | 161030008 | 0.22 | 85 | 4.36E-04 | 9.15E-07 | 1.20E-08 | 4.54E-10 | 3.21E-06 | 1.11E-06 | 2.62E-09 |
| 6 | 161028009 | 161032008 | 0.34 | 88 | 7.43E-03 | 2.03E-05 | 1.44E-15 | 7.42E-12 | 2.62E-09 | 5.77E-09 | 8.66E-15 |
| 6 | 161030009 | 161034008 | 0.47 | 95 | 7.74E-02 | 2.58E-03 | 3.01E-28 | 3.55E-32 | 1.45E-23 | 8.46E-24 | 2.13E-31 |
| 6 | 161032009 | 161036008 | 0.19 | 44 | 7.10E-01 | 4.21E-01 | 1.26E-12 | 1.80E-24 | 6.05E-24 | 3.18E-24 | 5.80E-24 |
| 6 | 161034009 | 161038008 | 0.36 | 20 | 1.27E-01 | 1.21E-01 | 7.39E-08 | 1.43E-10 | 3.71E-06 | 3.25E-06 | 8.55E-10 |
| 6 | 161036009 | 161040008 | 0.36 | 20 | 1.27E-01 | 1.21E-01 | 7.39E-08 | 1.43E-10 | 3.71E-06 | 3.25E-06 | 8.55E-10 |
| 6 | 161050009 | 161054008 | 0.15 | 14 | 9.12E-05 | 1.08E-07 | 4.87E-14 | 3.09E-14 | 1.32E-11 | 1.02E-11 | 1.13E-13 |
| 6 | 161052009 | 161056008 | 0.35 | 49 | 1.88E-04 | 7.35E-07 | 5.20E-23 | 4.25E-13 | 2.86E-11 | 2.22E-11 | 1.11E-16 |
| 6 | 161054009 | 161058008 | 0.31 | 54 | 9.95E-02 | 3.70E-03 | 3.24E-13 | 1.63E-09 | 4.96E-06 | 2.69E-06 | 1.94E-12 |
| 6 | 161068009 | 161072008 | 0.50 | 92 | 5.69E-10 | 1.06E-12 | 1.72E-13 | 1.75E-11 | 7.07E-08 | 9.04E-08 | 8.79E-13 |
| 6 | 161070009 | 161074008 | 0.44 | 54 | 1.69E-09 | 1.70E-09 | 6.45E-19 | 1.85E-08 | 7.45E-10 | 1.33E-09 | 1.11E-16 |
| 6 | 161072009 | 161076008 | 0.38 | 42 | 1.39E-02 | 8.73E-05 | 2.22E-15 | 2.13E-07 | 6.30E-10 | 1.15E-09 | 1.33E-14 |
| 6 | 161074009 | 161078008 | 0.28 | 52 | 3.34E-03 | 6.69E-08 | 1.81E-19 | 1.51E-13 | 1.14E-06 | 9.62E-07 | 1.11E-16 |
| 6 | 161076009 | 161080008 | 0.24 | 62 | 1.18E-02 | 2.19E-03 | 2.21E-23 | 1.16E-28 | 1.28E-29 | 1.15E-29 | 3.45E-29 |
| 6 | 161078009 | 161082008 | 0.14 | 53 | 4.88E-01 | 7.16E-01 | 1.61E-27 | 4.87E-32 | 8.21E-30 | 6.10E-30 | 2.88E-31 |
| 6 | 161082009 | 161086008 | 0.55 | 51 | 3.08E-01 | 3.78E-01 | 1.00E-10 | 2.57E-10 | 4.66E-06 | 3.70E-06 | 4.33E-10 |
| 6 | 161084009 | 161088008 | 0.57 | 61 | 4.14E-01 | 3.22E-01 | 1.64E-12 | 6.12E-13 | 1.20E-30 | 5.47E-31 | 2.25E-30 |
| 6 | 161086009 | 161090008 | 0.49 | 74 | 6.42E-01 | 1.48E-01 | 5.69E-10 | 1.15E-10 | 1.36E-30 | 4.53E-31 | 2.04E-30 |
| 6 | 161088009 | 161092008 | 0.45 | 65 | 2.24E-01 | 8.40E-02 | 2.51E-13 | 1.09E-09 | 3.15E-22 | 2.00E-22 | 7.35E-22 |
| 6 | 161090009 | 161094008 | 0.30 | 61 | 2.07E-06 | 3.33E-07 | 9.67E-55 | 3.56E-45 | 1.26E-29 | 9.76E-30 | 5.80E-54 |
| 6 | 161092009 | 161096008 | 0.31 | 58 | 2.81E-01 | 3.54E-02 | 6.74E-32 | 3.94E-23 | 1.21E-29 | 1.02E-29 | 4.00E-31 |
| 6 | 161098009 | 161102008 | 0.28 | 60 | 2.49E-01 | 2.30E-01 | 1.77E-10 | 2.08E-07 | 8.87E-07 | 8.53E-07 | 1.06E-09 |
| 6 | 161100009 | 161104008 | 0.34 | 56 | 5.59E-01 | 3.02E-01 | 1.59E-11 | 4.83E-07 | 4.91E-05 | 7.55E-05 | 9.53E-11 |
| 6 | 161114009 | 161118008 | 0.35 | 61 | 9.12E-01 | 5.55E-01 | 2.15E-08 | 1.80E-07 | 3.63E-05 | 3.32E-05 | 1.15E-07 |
| 6 | 161118009 | 161122008 | 0.49 | 78 | 3.89E-01 | 6.25E-01 | 2.38E-09 | 9.33E-07 | 2.26E-22 | 2.51E-22 | 7.14E-22 |
| 6 | 161120009 | 161124008 | 0.50 | 71 | 6.92E-04 | 4.26E-02 | 1.35E-13 | 3.15E-06 | 2.08E-22 | 2.61E-22 | 6.94E-22 |
| 6 | 161132009 | 161136008 | 0.34 | 59 | 2.06E-01 | 2.32E-01 | 2.04E-11 | 3.80E-07 | 1.25E-12 | 1.67E-12 | 4.15E-12 |
| 6 | 161134009 | 161138008 | 0.26 | 61 | 4.97E-01 | 7.88E-01 | 4.22E-13 | 1.12E-07 | 1.19E-12 | 1.16E-12 | 1.47E-12 |

| | | | | | | | | | | | |
|---|---|---|---|---|---|---|---|---|---|---|---|
| 6 | 161150009 | 161154008 | 0.29 | 56 | 3.43E-03 | 1.93E-03 | 7.01E-07 | 3.98E-05 | 1.29E-09 | 1.08E-09 | 3.53E-09 |
| 6 | 161152009 | 161156008 | 0.40 | 61 | 5.23E-01 | 3.18E-01 | 2.45E-06 | 1.53E-05 | 1.73E-09 | 1.47E-09 | 4.76E-09 |
| 6 | 161174009 | 161178008 | 0.20 | 50 | 3.74E-01 | 3.05E-01 | 2.62E-11 | 7.68E-12 | 4.01E-11 | 2.77E-11 | 2.61E-11 |
| 6 | 161176009 | 161180008 | 0.17 | 48 | 8.34E-01 | 3.49E-01 | 6.84E-12 | 3.85E-07 | 2.46E-11 | 2.18E-11 | 2.58E-11 |
| 6 | 161178009 | 161182008 | 0.42 | 64 | 3.03E-01 | 2.24E-01 | 1.33E-15 | 1.48E-05 | 4.64E-11 | 4.70E-11 | 8.22E-15 |
| 6 | 161180009 | 161184008 | 0.42 | 52 | 8.06E-01 | 7.19E-01 | 2.74E-14 | 6.11E-09 | 5.50E-11 | 4.79E-11 | 1.65E-13 |
| 6 | 161194009 | 161198008 | 0.59 | 68 | 1.03E-01 | 1.98E-01 | 1.69E-07 | 8.10E-04 | 1.97E-10 | 2.67E-10 | 6.80E-10 |
| 6 | 161196009 | 161200008 | 0.53 | 74 | 4.99E-01 | 4.04E-01 | 5.17E-07 | 7.16E-04 | 1.81E-10 | 2.40E-10 | 6.20E-10 |
| 6 | 161202009 | 161206008 | 0.23 | 54 | 1.34E-03 | 1.40E-03 | 2.63E-07 | 2.87E-05 | 2.01E-10 | 2.25E-10 | 6.37E-10 |
| 6 | 161204009 | 161208008 | 0.25 | 59 | 1.88E-03 | 2.87E-03 | 1.17E-07 | 1.61E-04 | 2.59E-10 | 2.50E-10 | 7.63E-10 |
| 6 | 161212009 | 161216008 | 0.27 | 44 | 9.30E-01 | 6.87E-01 | 1.04E-08 | 2.69E-08 | 4.39E-08 | 3.01E-08 | 3.17E-08 |
| 6 | 161254009 | 161258008 | 0.26 | 40 | 4.17E-01 | 2.45E-01 | 8.00E-08 | 1.89E-08 | 8.45E-08 | 7.45E-08 | 6.62E-08 |
| 6 | 161278009 | 161282008 | 0.58 | 76 | 8.44E-01 | 7.32E-01 | 1.36E-07 | 3.03E-08 | 4.51E-08 | 4.46E-08 | 7.07E-08 |
| 6 | 161288009 | 161292008 | 0.49 | 66 | 3.74E-02 | 1.66E-02 | 3.73E-05 | 6.65E-03 | 4.40E-09 | 6.14E-09 | 1.54E-08 |
| 6 | 161290009 | 161294008 | 0.36 | 65 | 2.85E-01 | 5.41E-01 | 1.87E-09 | 4.20E-04 | 3.07E-09 | 4.44E-09 | 5.53E-09 |
| 6 | 161302009 | 161306008 | 0.35 | 53 | 8.98E-01 | 9.47E-01 | 8.49E-10 | 1.79E-04 | 5.49E-09 | 8.63E-09 | 4.07E-09 |
| 6 | 161304009 | 161308008 | 0.25 | 53 | 1.23E-01 | 1.39E-01 | 3.09E-08 | 9.51E-04 | 4.07E-09 | 5.65E-09 | 1.32E-08 |
| 6 | 161334009 | 161338008 | 0.58 | 54 | 6.15E-02 | 4.34E-02 | 1.29E-04 | 1.63E-02 | 2.62E-10 | 4.95E-10 | 1.03E-09 |
| 6 | 161336009 | 161340008 | 0.88 | 74 | 2.50E-03 | 4.81E-03 | 2.60E-04 | 1.44E-02 | 3.30E-10 | 7.39E-10 | 1.37E-09 |
| 6 | 161340009 | 161344008 | 0.50 | 56 | 2.32E-01 | 1.65E-01 | 1.02E-03 | 2.11E-02 | 1.03E-10 | 1.91E-10 | 4.02E-10 |
| 6 | 161342009 | 161346008 | 0.44 | 59 | 1.46E-02 | 2.08E-02 | 2.34E-04 | 1.46E-02 | 9.11E-11 | 1.68E-10 | 3.54E-10 |

cMAF = cumulative minor allele frequency

**Table S3**

Significant 4kb sliding windows for Lp(a) levels in EAs. The p-value threshold for genome-wide significance is $3.75 \times 10^{-8}$. Seven methods are compared: ACAT-V(1,1), ACAT-V(1,25), SKAT(1,1), SKAT(1,25), Burden(1,1), Burden(1,25) and the omnibus test ACAT-O that combines the other six tests, where the two numbers in the parentheses indicate the choice of the beta(MAF) weight parameters $a_1$ and $a_2$ in the test.

| Chr. | Start position (bp) | End position (bp) | cMAF | #SNV | P-values | | | | | | |
|---|---|---|---|---|---|---|---|---|---|---|---|
| | | | | | Burden (1,25) | Burden (1,1) | SKAT (1,25) | SKAT (1,1) | ACAT-V (1,25) | ACAT-V (1,1) | ACAT-O |
| 6 | 160596009 | 160600008 | 0.04 | 34 | 5.10E-03 | 3.04E-04 | 6.78E-09 | 4.55E-09 | 4.45E-09 | 3.72E-09 | 6.97E-09 |
| 6 | 160598009 | 160602008 | 0.07 | 43 | 1.91E-03 | 2.13E-04 | 1.03E-08 | 4.66E-09 | 9.67E-09 | 7.40E-09 | 1.09E-08 |
| 6 | 160608009 | 160612008 | 0.22 | 51 | 1.28E-03 | 5.17E-03 | 1.17E-03 | 1.25E-02 | 3.74E-08 | 7.17E-08 | 1.48E-07 |
| 6 | 160748009 | 160752008 | 0.06 | 32 | 2.44E-05 | 1.03E-06 | 7.94E-11 | 1.06E-10 | 2.10E-11 | 1.86E-11 | 4.86E-11 |
| 6 | 160750009 | 160754008 | 0.09 | 33 | 1.36E-03 | 6.22E-03 | 3.52E-11 | 2.36E-08 | 1.50E-11 | 2.72E-11 | 4.55E-11 |
| 6 | 160764009 | 160768008 | 0.11 | 38 | 1.35E-05 | 4.62E-07 | 7.04E-10 | 1.90E-08 | 3.57E-08 | 4.80E-08 | 3.94E-09 |
| 6 | 160766009 | 160770008 | 0.09 | 40 | 6.54E-02 | 8.28E-03 | 2.86E-07 | 2.38E-08 | 4.62E-08 | 3.19E-08 | 6.09E-08 |
| 6 | 160814009 | 160818008 | 0.04 | 34 | 1.16E-05 | 1.11E-06 | 7.39E-09 | 1.18E-08 | 5.01E-08 | 4.27E-08 | 2.27E-08 |
| 6 | 160816009 | 160820008 | 0.05 | 35 | 6.53E-03 | 1.57E-03 | 3.03E-07 | 2.68E-08 | 1.06E-07 | 8.11E-08 | 9.60E-08 |
| 6 | 160826009 | 160830008 | 0.02 | 23 | 6.84E-02 | 3.47E-02 | 1.70E-08 | 9.39E-09 | 6.01E-09 | 5.80E-09 | 1.19E-08 |
| 6 | 160828009 | 160832008 | 0.07 | 34 | 3.41E-03 | 3.13E-03 | 6.82E-04 | 5.26E-04 | 3.75E-08 | 3.58E-08 | 1.10E-07 |
| 6 | 160836009 | 160840008 | 0.07 | 29 | 1.44E-05 | 5.20E-06 | 1.10E-10 | 7.02E-10 | 2.14E-06 | 1.70E-06 | 5.71E-10 |
| 6 | 160846009 | 160850008 | 0.18 | 43 | 5.21E-01 | 8.16E-01 | 3.19E-08 | 2.93E-09 | 1.66E-10 | 2.05E-10 | 5.32E-10 |
| 6 | 160848009 | 160852008 | 0.08 | 29 | 4.86E-04 | 2.38E-05 | 1.43E-12 | 1.83E-10 | 9.00E-11 | 8.78E-11 | 8.24E-12 |
| 6 | 160850009 | 160854008 | 0.08 | 29 | 5.86E-13 | 1.19E-14 | 5.24E-23 | 4.40E-14 | 4.04E-11 | 3.50E-11 | 1.11E-16 |
| 6 | 160852009 | 160856008 | 0.11 | 39 | 8.65E-06 | 2.55E-05 | 2.28E-23 | 3.95E-12 | 4.80E-11 | 4.27E-11 | 1.11E-16 |
| 6 | 160864009 | 160868008 | 0.05 | 46 | 2.97E-01 | 2.24E-01 | 1.51E-07 | 1.70E-08 | 2.60E-07 | 2.20E-07 | 8.14E-08 |
| 6 | 160894009 | 160898008 | 0.09 | 22 | 7.44E-01 | 6.33E-01 | 6.30E-09 | 5.34E-07 | 9.01E-08 | 1.17E-07 | 3.33E-08 |
| 6 | 160908009 | 160912008 | 0.21 | 68 | 7.59E-01 | 2.92E-01 | 2.05E-05 | 9.87E-07 | 7.20E-09 | 8.25E-09 | 2.30E-08 |
| 6 | 160910009 | 160914008 | 0.23 | 53 | 3.77E-01 | 1.61E-01 | 8.55E-04 | 1.12E-04 | 7.95E-09 | 9.05E-09 | 2.54E-08 |
| 6 | 160914009 | 160918008 | 0.17 | 74 | 9.09E-01 | 8.98E-01 | 2.00E-06 | 1.86E-04 | 3.60E-09 | 3.98E-09 | 1.13E-08 |
| 6 | 160916009 | 160920008 | 0.17 | 78 | 7.08E-01 | 6.25E-01 | 4.04E-05 | 2.54E-05 | 3.69E-09 | 3.84E-09 | 1.13E-08 |
| 6 | 160920009 | 160924008 | 0.16 | 51 | 5.25E-01 | 3.35E-01 | 6.51E-04 | 3.54E-04 | 9.37E-09 | 1.05E-08 | 2.97E-08 |
| 6 | 160922009 | 160926008 | 0.11 | 59 | 5.99E-01 | 5.03E-01 | 1.40E-04 | 1.14E-04 | 6.85E-09 | 6.39E-09 | 1.98E-08 |
| 6 | 160940009 | 160944008 | 0.14 | 38 | 3.94E-01 | 4.40E-02 | 1.47E-06 | 6.42E-10 | 6.69E-08 | 2.17E-08 | 3.70E-09 |
| 6 | 160942009 | 160946008 | 0.11 | 33 | 1.55E-01 | 1.05E-02 | 3.28E-08 | 4.32E-10 | 3.79E-08 | 1.61E-08 | 2.46E-09 |
| 6 | 160950009 | 160954008 | 0.21 | 39 | 6.38E-01 | 4.19E-01 | 1.80E-13 | 1.35E-11 | 1.09E-06 | 3.32E-07 | 1.07E-12 |
| 6 | 160958009 | 160962008 | 0.03 | 33 | 1.37E-05 | 1.28E-06 | 5.01E-09 | 5.29E-09 | 8.97E-09 | 7.78E-09 | 9.53E-09 |
| 6 | 160960009 | 160964008 | 0.09 | 37 | 1.46E-02 | 2.02E-02 | 1.21E-04 | 9.19E-04 | 2.97E-08 | 3.26E-08 | 9.33E-08 |
| 6 | 161004009 | 161008008 | 0.09 | 39 | 8.22E-03 | 5.74E-05 | 1.22E-09 | 1.08E-09 | 1.12E-08 | 5.14E-09 | 2.95E-09 |
| 6 | 161006009 | 161010008 | 0.13 | 48 | 1.66E-04 | 3.25E-07 | 8.03E-12 | 1.29E-07 | 8.75E-09 | 7.22E-09 | 4.81E-11 |
| 6 | 161008009 | 161012008 | 0.16 | 46 | 1.86E-02 | 1.27E-02 | 1.05E-10 | 1.69E-08 | 2.57E-08 | 1.72E-08 | 6.20E-10 |
| 6 | 161010009 | 161014008 | 0.12 | 44 | 4.88E-10 | 7.67E-14 | 2.22E-27 | 1.33E-15 | 7.05E-23 | 1.10E-22 | 1.33E-26 |
| 6 | 161012009 | 161016008 | 0.09 | 43 | 7.26E-01 | 8.40E-01 | 7.95E-13 | 1.24E-08 | 5.66E-23 | 7.55E-23 | 1.94E-22 |
| 6 | 161014009 | 161018008 | 0.12 | 41 | 7.68E-01 | 1.58E-01 | 1.63E-10 | 1.79E-12 | 1.70E-10 | 7.85E-11 | 1.03E-11 |
| 6 | 161016009 | 161020008 | 0.10 | 39 | 4.68E-03 | 2.78E-05 | 4.09E-09 | 6.00E-11 | 1.48E-10 | 6.87E-11 | 1.57E-10 |
| 6 | 161020009 | 161024008 | 0.10 | 40 | 4.11E-01 | 9.28E-02 | 7.82E-09 | 2.31E-08 | 6.39E-06 | 4.01E-06 | 3.50E-08 |
| 6 | 161030009 | 161034008 | 0.13 | 68 | 6.77E-08 | 1.55E-08 | 1.79E-14 | 2.11E-15 | 5.99E-11 | 6.81E-11 | 1.15E-14 |
| 6 | 161032009 | 161036008 | 0.09 | 31 | 7.25E-11 | 4.46E-11 | 3.31E-14 | 4.22E-15 | 4.71E-11 | 5.51E-11 | 2.26E-14 |
| 6 | 161060009 | 161064008 | 0.11 | 9 | 1.95E-03 | 4.34E-05 | 1.53E-10 | 8.73E-11 | 2.77E-10 | 1.62E-10 | 2.16E-10 |
| 6 | 161062009 | 161066008 | 0.11 | 10 | 4.44E-03 | 6.72E-05 | 1.59E-10 | 8.76E-11 | 2.76E-10 | 1.62E-10 | 2.18E-10 |
| 6 | 161084009 | 161088008 | 0.06 | 36 | 5.41E-03 | 1.05E-03 | 4.08E-11 | 7.03E-13 | 3.55E-13 | 2.94E-13 | 7.83E-13 |
| 6 | 161086009 | 161090008 | 0.23 | 41 | 5.05E-03 | 1.39E-02 | 9.57E-06 | 5.65E-03 | 1.02E-12 | 1.60E-12 | 3.73E-12 |
| 6 | 161108009 | 161112008 | 0.26 | 48 | 1.62E-01 | 3.52E-01 | 1.52E-09 | 4.40E-05 | 3.72E-23 | 6.26E-23 | 1.40E-22 |
| 6 | 161110009 | 161114008 | 0.12 | 49 | 4.11E-04 | 4.48E-04 | 1.90E-23 | 8.84E-08 | 1.64E-23 | 2.55E-23 | 3.92E-23 |
| 6 | 161170009 | 161174008 | 0.19 | 51 | 1.47E-02 | 1.39E-02 | 1.72E-05 | 1.99E-06 | 7.79E-11 | 7.17E-11 | 2.24E-10 |
| 6 | 161172009 | 161176008 | 0.18 | 49 | 5.36E-04 | 4.03E-04 | 3.49E-06 | 5.23E-07 | 7.10E-11 | 6.47E-11 | 2.03E-10 |
| 6 | 161174009 | 161178008 | 0.12 | 39 | 3.01E-07 | 2.97E-08 | 1.31E-09 | 6.74E-07 | 3.64E-09 | 4.92E-09 | 4.69E-09 |
| 6 | 161176009 | 161180008 | 0.11 | 39 | 1.86E-05 | 1.64E-06 | 1.47E-09 | 2.41E-06 | 3.83E-09 | 4.98E-09 | 5.24E-09 |
| 6 | 161282009 | 161286008 | 0.14 | 32 | 1.09E-01 | 6.92E-02 | 2.12E-09 | 2.00E-05 | 2.02E-07 | 2.33E-07 | 1.25E-08 |
| 6 | 161380009 | 161384008 | 0.13 | 25 | 4.77E-01 | 5.14E-01 | 1.82E-11 | 6.50E-05 | 3.72E-12 | 6.24E-12 | 1.24E-11 |
| 6 | 161382009 | 161386008 | 0.13 | 26 | 8.13E-01 | 8.59E-01 | 5.51E-07 | 4.17E-04 | 4.38E-12 | 6.19E-12 | 1.54E-11 |
| 6 | 161442009 | 161446008 | 0.08 | 28 | 3.66E-02 | 4.18E-02 | 3.90E-08 | 6.67E-03 | 2.35E-08 | 6.94E-08 | 7.27E-08 |

cMAF = cumulative minor allele frequency

**Table S4**

Significant 4kb sliding windows for neutrophil count levels in AAs. The p-value threshold for genome-wide significance is $3.75 \times 10^{-8}$. Seven methods are compared: ACAT-V(1,1), ACAT-V(1,25), SKAT(1,1), SKAT(1,25), Burden(1,1), Burden(1,25) and the omnibus test ACAT-O that combines the other six tests, where the two numbers in the parentheses indicate the choice of the beta(MAF) weight parameters $a_1$ and $a_2$ in the test.

| Chr. | Start position (bp) | End position (bp) | cMAF | #SNV | P-values | | | | | | |
|---|---|---|---|---|---|---|---|---|---|---|---|
| | | | | | Burden (1,25) | Burden (1,1) | SKAT (1,25) | SKAT (1,1) | ACAT-V (1,25) | ACAT-V (1,1) | ACAT-O |
| 1 | 118642150 | 118646149 | 0.19 | 50 | 6.61E-01 | 5.87E-01 | 3.86E-04 | 1.38E-08 | 2.05E-06 | 4.85E-07 | 7.99E-08 |
| 1 | 154634150 | 154638149 | 0.44 | 44 | 6.61E-02 | 6.05E-04 | 2.47E-06 | 6.79E-09 | 6.46E-06 | 2.32E-06 | 4.04E-08 |
| 1 | 154828150 | 154832149 | 0.45 | 81 | 7.74E-01 | 8.96E-01 | 4.89E-03 | 2.66E-04 | 4.21E-09 | 1.85E-09 | 7.71E-09 |
| 1 | 154830150 | 154834149 | 0.24 | 56 | 8.65E-01 | 3.24E-01 | 5.19E-05 | 1.09E-07 | 2.12E-09 | 9.52E-10 | 3.92E-09 |
| 1 | 155052150 | 155056149 | 0.17 | 38 | 1.10E-02 | 8.95E-03 | 3.74E-08 | 2.77E-07 | 7.13E-05 | 5.28E-05 | 1.98E-07 |
| 1 | 155062150 | 155066149 | 0.28 | 59 | 1.37E-01 | 5.01E-02 | 1.74E-07 | 2.05E-10 | 9.82E-07 | 3.05E-07 | 1.23E-09 |
| 1 | 155064150 | 155068149 | 0.12 | 38 | 5.51E-01 | 4.08E-02 | 1.21E-06 | 8.36E-09 | 4.41E-07 | 1.16E-07 | 4.57E-08 |
| 1 | 156114150 | 156118149 | 0.22 | 44 | 3.37E-01 | 5.15E-03 | 1.75E-04 | 3.18E-09 | 5.92E-06 | 1.74E-06 | 1.90E-08 |
| 1 | 156116150 | 156120149 | 0.20 | 49 | 4.13E-01 | 2.25E-03 | 1.11E-04 | 1.65E-09 | 5.75E-06 | 1.48E-06 | 9.88E-09 |
| 1 | 156744150 | 156748149 | 0.25 | 62 | 3.66E-01 | 8.51E-01 | 5.31E-07 | 1.01E-10 | 6.86E-10 | 2.29E-10 | 3.82E-10 |
| 1 | 156746150 | 156750149 | 0.31 | 60 | 3.20E-01 | 7.31E-03 | 4.85E-09 | 1.22E-13 | 6.63E-10 | 2.99E-10 | 7.29E-13 |
| 1 | 156756150 | 156760149 | 0.19 | 42 | 4.92E-01 | 1.98E-02 | 2.92E-05 | 2.84E-10 | 8.18E-10 | 2.39E-10 | 6.72E-10 |
| 1 | 156758150 | 156762149 | 0.17 | 44 | 9.70E-01 | 8.55E-02 | 4.42E-06 | 2.80E-10 | 6.50E-10 | 2.17E-10 | 6.17E-10 |
| 1 | 156784150 | 156788149 | 0.20 | 59 | 4.35E-01 | 2.62E-02 | 4.70E-04 | 1.26E-10 | 1.13E-09 | 3.03E-10 | 4.96E-10 |
| 1 | 156786150 | 156790149 | 0.34 | 52 | 9.77E-01 | 2.52E-01 | 3.30E-03 | 1.31E-05 | 1.56E-09 | 5.61E-10 | 2.47E-09 |
| 1 | 156906150 | 156910149 | 0.26 | 66 | 7.66E-01 | 3.70E-01 | 3.38E-03 | 2.43E-08 | 1.96E-07 | 4.91E-08 | 9.00E-08 |
| 1 | 157074150 | 157078149 | 0.30 | 66 | 5.25E-01 | 9.19E-02 | 9.64E-05 | 1.38E-07 | 1.98E-08 | 6.84E-09 | 2.94E-08 |
| 1 | 157076150 | 157080149 | 0.46 | 86 | 9.21E-01 | 8.40E-01 | 3.52E-05 | 2.71E-08 | 2.90E-08 | 1.07E-08 | 3.63E-08 |
| 1 | 157396150 | 157400149 | 0.39 | 54 | 9.27E-01 | 7.95E-01 | 1.97E-06 | 4.15E-08 | 1.49E-08 | 5.05E-09 | 2.07E-08 |
| 1 | 157398150 | 157402149 | 0.42 | 71 | 6.26E-01 | 7.35E-01 | 1.06E-04 | 6.68E-08 | 1.49E-08 | 5.37E-09 | 2.24E-08 |
| 1 | 157400150 | 157404149 | 0.28 | 74 | 2.84E-01 | 8.14E-02 | 1.88E-05 | 6.48E-10 | 3.47E-08 | 9.50E-09 | 3.58E-09 |
| 1 | 157402150 | 157406149 | 0.31 | 62 | 9.94E-02 | 4.95E-01 | 1.39E-05 | 6.18E-10 | 3.95E-08 | 1.11E-08 | 3.46E-09 |
| 1 | 157412150 | 157416149 | 0.45 | 74 | 2.04E-01 | 7.24E-01 | 1.18E-04 | 6.52E-07 | 6.48E-09 | 2.51E-09 | 1.08E-08 |
| 1 | 157414150 | 157418149 | 0.41 | 65 | 4.99E-02 | 1.59E-01 | 8.79E-05 | 9.07E-06 | 6.08E-09 | 2.32E-09 | 1.01E-08 |
| 1 | 157450150 | 157454149 | 0.37 | 56 | 8.73E-01 | 8.30E-01 | 1.81E-04 | 1.37E-06 | 7.28E-08 | 3.72E-08 | 1.45E-07 |
| 1 | 157452150 | 157456149 | 0.28 | 59 | 7.70E-01 | 7.83E-01 | 6.41E-06 | 4.39E-09 | 7.29E-08 | 2.72E-08 | 2.15E-08 |
| 1 | 157552150 | 157556149 | 0.50 | 65 | 6.21E-01 | 3.15E-01 | 1.22E-03 | 2.97E-08 | 4.72E-08 | 1.37E-08 | 4.70E-08 |
| 1 | 157554150 | 157558149 | 0.43 | 61 | 1.03E-01 | 3.11E-02 | 4.32E-04 | 3.38E-08 | 3.88E-08 | 1.17E-08 | 4.26E-08 |
| 1 | 157564150 | 157568149 | 0.30 | 59 | 1.89E-01 | 3.75E-01 | 1.91E-03 | 8.80E-06 | 8.24E-08 | 2.64E-08 | 1.20E-07 |
| 1 | 157670150 | 157674149 | 0.24 | 48 | 6.89E-01 | 2.31E-01 | 1.22E-06 | 1.03E-11 | 1.62E-07 | 4.76E-08 | 6.15E-11 |
| 1 | 157672150 | 157676149 | 0.23 | 47 | 2.36E-02 | 7.59E-05 | 1.32E-06 | 4.72E-11 | 1.61E-07 | 4.33E-08 | 2.83E-10 |
| 1 | 157924150 | 157928149 | 0.80 | 91 | 1.19E-01 | 1.12E-01 | 7.63E-02 | 2.17E-03 | 5.75E-10 | 1.40E-10 | 6.76E-10 |
| 1 | 157926150 | 157930149 | 0.75 | 88 | 9.24E-02 | 1.23E-01 | 4.88E-02 | 1.05E-03 | 5.67E-10 | 1.33E-10 | 6.46E-10 |
| 1 | 157956150 | 157960149 | 0.26 | 66 | 3.70E-01 | 8.78E-01 | 1.91E-05 | 1.35E-09 | 3.57E-10 | 1.40E-10 | 5.60E-10 |
| 1 | 157958150 | 157962149 | 0.25 | 66 | 4.86E-01 | 8.54E-01 | 3.02E-04 | 1.61E-08 | 3.32E-10 | 1.32E-10 | 5.64E-10 |
| 1 | 158572150 | 158576149 | 0.28 | 46 | 1.79E-04 | 1.08E-04 | 1.57E-08 | 6.37E-09 | 2.43E-07 | 2.56E-07 | 2.62E-08 |
| 1 | 158728150 | 158732149 | 0.30 | 63 | 8.77E-01 | 6.36E-01 | 1.62E-03 | 1.18E-11 | 6.56E-10 | 1.41E-10 | 6.42E-11 |
| 1 | 158730150 | 158734149 | 0.46 | 48 | 3.32E-01 | 1.63E-01 | 3.11E-08 | 2.45E-13 | 7.42E-10 | 2.26E-10 | 1.47E-12 |
| 1 | 158764150 | 158768149 | 0.23 | 39 | 2.40E-04 | 6.01E-04 | 8.78E-12 | 6.05E-10 | 3.72E-05 | 2.65E-05 | 5.19E-11 |
| 1 | 158778150 | 158782149 | 0.42 | 79 | 6.97E-02 | 5.38E-02 | 4.38E-06 | 2.95E-08 | 6.94E-08 | 6.38E-08 | 9.35E-08 |
| 1 | 158894150 | 158898149 | 0.25 | 46 | 8.78E-01 | 2.79E-01 | 2.50E-06 | 1.50E-08 | 3.82E-06 | 2.81E-06 | 8.88E-08 |
| 1 | 159166150 | 159170149 | 0.32 | 57 | 3.80E-03 | 1.12E-05 | 1.18E-09 | 8.55E-09 | 1.62E-06 | 6.16E-07 | 6.19E-09 |
| 1 | 159168150 | 159172149 | 0.41 | 62 | 2.17E-04 | 1.11E-05 | 2.80E-11 | 4.26E-10 | 8.53E-05 | 2.71E-05 | 1.58E-10 |
| 1 | 159170150 | 159174149 | 0.43 | 64 | 8.54E-06 | 4.82E-07 | 1.68E-09 | 3.76E-09 | 1.10E-07 | 6.61E-08 | 6.77E-09 |
| 1 | 159172150 | 159176149 | 0.31 | 57 | 8.31E-08 | 4.50E-10 | 2.52E-10 | 1.61E-10 | 3.47E-08 | 2.40E-08 | 4.81E-10 |
| 1 | 159174150 | 159178149 | 0.27 | 56 | 7.41E-12 | 4.66E-19 | 3.68E-11 | 4.75E-22 | 3.72E-13 | 1.24E-13 | 2.19E-21 |
| 1 | 159176150 | 159180149 | 0.32 | 53 | 8.29E-08 | 2.08E-12 | 6.91E-09 | 2.87E-20 | 4.61E-13 | 1.51E-13 | 1.11E-16 |
| 1 | 159200150 | 159204149 | 0.32 | 48 | 5.25E-01 | 3.35E-01 | 7.43E-05 | 8.61E-10 | 9.04E-11 | 2.54E-11 | 1.16E-10 |
| 1 | 159202150 | 159206149 | 0.33 | 56 | 1.08E-01 | 8.47E-03 | 3.38E-07 | 1.43E-12 | 8.24E-11 | 2.60E-11 | 8.00E-12 |
| 1 | 159220150 | 159224149 | 0.23 | 49 | 9.60E-02 | 6.24E-04 | 7.81E-08 | 1.15E-09 | 3.15E-09 | 9.77E-10 | 2.70E-09 |
| 1 | 159222150 | 159226149 | 0.27 | 48 | 3.21E-05 | 3.25E-07 | 8.76E-05 | 2.52E-10 | 3.76E-09 | 1.08E-09 | 1.16E-09 |
| 1 | 159230150 | 159234149 | 0.20 | 47 | 8.81E-03 | 3.09E-04 | 6.63E-06 | 1.53E-09 | 6.33E-08 | 2.97E-08 | 8.55E-09 |
| 1 | 159288150 | 159292149 | 0.25 | 42 | 7.70E-02 | 9.25E-04 | 2.22E-10 | 2.44E-15 | 3.86E-08 | 1.71E-08 | 1.49E-14 |
| 1 | 159290150 | 159294149 | 0.18 | 38 | 1.12E-07 | 1.83E-11 | 3.65E-14 | 7.45E-18 | 2.60E-08 | 1.20E-08 | 1.11E-16 |
| 1 | 159292150 | 159296149 | 0.22 | 55 | 7.34E-04 | 7.86E-07 | 1.89E-05 | 3.61E-11 | 4.56E-08 | 1.17E-08 | 2.15E-10 |
| 1 | 159294150 | 159298149 | 0.50 | 88 | 9.63E-02 | 9.94E-03 | 2.32E-03 | 4.51E-06 | 1.09E-07 | 2.82E-08 | 1.34E-07 |
| 1 | 159308150 | 159312149 | 0.24 | 51 | 2.05E-02 | 6.14E-03 | 4.44E-05 | 2.56E-08 | 1.92E-07 | 5.65E-08 | 9.68E-08 |
| 1 | 159310150 | 159314149 | 0.22 | 48 | 3.96E-04 | 1.13E-05 | 4.75E-07 | 4.04E-09 | 1.86E-07 | 5.12E-08 | 2.19E-08 |
| 1 | 159314150 | 159318149 | 0.25 | 60 | 3.58E-03 | 8.96E-06 | 2.79E-06 | 3.90E-11 | 1.66E-07 | 3.92E-08 | 2.34E-10 |
| 1 | 159316150 | 159320149 | 0.27 | 74 | 3.73E-07 | 1.28E-10 | 3.03E-06 | 5.01E-10 | 6.20E-08 | 1.63E-08 | 6.08E-10 |
| 1 | 159318150 | 159322149 | 0.19 | 70 | 8.54E-04 | 1.11E-05 | 1.89E-04 | 8.46E-09 | 7.25E-08 | 1.86E-08 | 3.23E-08 |
| 1 | 159322150 | 159326149 | 0.13 | 54 | 4.70E-01 | 1.40E-02 | 3.15E-04 | 9.96E-09 | 5.16E-08 | 1.26E-08 | 3.02E-08 |
| 1 | 159324150 | 159328149 | 0.23 | 45 | 2.53E-02 | 6.92E-06 | 1.50E-08 | 4.66E-15 | 1.21E-08 | 5.31E-09 | 2.81E-14 |
| 1 | 159326150 | 159330149 | 0.25 | 45 | 6.24E-03 | 1.19E-05 | 1.78E-07 | 4.17E-14 | 1.24E-08 | 5.65E-09 | 2.51E-13 |

| | | | | | | | | | | | |
|---|---|---|---|---|---|---|---|---|---|---|---|
| 1 | 159328150 | 159332149 | 0.23 | 48 | 1.36E-01 | 5.36E-02 | 1.42E-05 | 1.19E-07 | 6.56E-08 | 2.10E-08 | 8.41E-08 |
| 1 | 159334150 | 159338149 | 0.35 | 66 | 1.64E-01 | 2.16E-02 | 3.90E-06 | 3.75E-11 | 3.43E-06 | 1.43E-06 | 2.25E-10 |
| 1 | 159336150 | 159340149 | 0.27 | 56 | 1.68E-02 | 4.59E-04 | 2.66E-08 | 5.06E-11 | 2.09E-05 | 1.06E-05 | 3.03E-10 |
| 1 | 159352150 | 159356149 | 0.38 | 62 | 4.59E-01 | 1.64E-01 | 3.96E-04 | 1.12E-08 | 1.34E-05 | 5.22E-06 | 6.69E-08 |
| 1 | 159354150 | 159358149 | 0.35 | 72 | 1.64E-01 | 3.91E-02 | 3.69E-04 | 6.30E-09 | 1.29E-05 | 4.61E-06 | 3.77E-08 |
| 1 | 159368150 | 159372149 | 0.25 | 50 | 4.60E-02 | 2.05E-02 | 7.24E-08 | 1.57E-08 | 2.10E-05 | 6.65E-06 | 7.73E-08 |
| 1 | 159370150 | 159374149 | 0.42 | 60 | 2.00E-02 | 9.13E-06 | 2.41E-11 | 1.08E-13 | 1.43E-07 | 6.88E-08 | 6.44E-13 |
| 1 | 159372150 | 159376149 | 0.58 | 70 | 7.06E-04 | 2.71E-06 | 5.72E-08 | 1.03E-09 | 1.62E-07 | 9.36E-08 | 5.98E-09 |
| 1 | 159384150 | 159388149 | 0.27 | 56 | 8.39E-02 | 5.89E-03 | 4.33E-08 | 2.05E-09 | 4.87E-07 | 4.01E-07 | 1.16E-08 |
| 1 | 159394150 | 159398149 | 0.27 | 53 | 2.78E-02 | 1.03E-02 | 8.97E-08 | 2.05E-07 | 1.39E-08 | 7.57E-09 | 2.73E-08 |
| 1 | 159396150 | 159400149 | 0.26 | 60 | 2.83E-04 | 8.37E-05 | 6.03E-08 | 3.14E-08 | 1.28E-08 | 7.41E-09 | 2.30E-08 |
| 1 | 159402150 | 159406149 | 0.26 | 57 | 2.04E-03 | 2.59E-05 | 7.59E-11 | 6.24E-10 | 7.39E-07 | 5.13E-07 | 4.06E-10 |
| 1 | 159404150 | 159408149 | 0.25 | 57 | 9.86E-06 | 4.96E-08 | 5.35E-08 | 1.99E-12 | 8.11E-07 | 5.62E-07 | 1.19E-11 |
| 1 | 159406150 | 159410149 | 0.38 | 57 | 3.50E-03 | 5.52E-04 | 6.67E-08 | 1.04E-08 | 7.04E-06 | 3.20E-06 | 5.36E-08 |
| 1 | 159408150 | 159412149 | 0.46 | 59 | 4.56E-03 | 2.18E-05 | 9.48E-11 | 1.51E-13 | 9.74E-10 | 3.47E-10 | 9.05E-13 |
| 1 | 159410150 | 159414149 | 0.41 | 53 | 1.43E-07 | 1.26E-10 | 6.79E-09 | 8.73E-10 | 7.31E-10 | 3.07E-10 | 4.33E-10 |
| 1 | 159416150 | 159420149 | 0.73 | 102 | 4.61E-05 | 5.97E-06 | 1.20E-09 | 2.30E-08 | 9.36E-06 | 5.53E-06 | 6.87E-09 |
| 1 | 159418150 | 159422149 | 0.75 | 98 | 9.09E-06 | 1.82E-06 | 9.54E-09 | 7.24E-08 | 9.60E-06 | 5.76E-06 | 5.02E-08 |
| 1 | 159428150 | 159432149 | 0.21 | 50 | 6.55E-02 | 3.03E-03 | 7.07E-06 | 2.54E-12 | 1.19E-09 | 2.81E-10 | 1.51E-11 |
| 1 | 159430150 | 159434149 | 0.21 | 57 | 2.97E-02 | 1.43E-03 | 1.32E-05 | 2.54E-12 | 9.94E-10 | 2.62E-10 | 1.51E-11 |
| 1 | 159436150 | 159440149 | 0.20 | 42 | 5.27E-02 | 8.45E-02 | 3.15E-08 | 4.57E-08 | 6.23E-06 | 3.14E-06 | 1.11E-07 |
| 1 | 159442150 | 159446149 | 0.29 | 61 | 8.08E-05 | 1.94E-07 | 4.62E-10 | 8.98E-11 | 9.19E-08 | 5.57E-08 | 4.50E-10 |
| 1 | 159444150 | 159448149 | 0.32 | 65 | 8.39E-08 | 1.43E-09 | 3.49E-11 | 9.08E-09 | 7.89E-08 | 5.57E-08 | 2.04E-10 |
| 1 | 159446150 | 159450149 | 0.23 | 62 | 6.06E-11 | 1.92E-13 | 3.23E-19 | 1.68E-09 | 1.65E-07 | 1.21E-07 | 1.11E-16 |
| 1 | 159448150 | 159452149 | 0.22 | 53 | 1.07E-05 | 3.20E-07 | 1.95E-10 | 4.66E-08 | 2.30E-06 | 2.86E-06 | 1.16E-09 |
| 1 | 159450150 | 159454149 | 0.42 | 50 | 1.71E-10 | 3.20E-10 | 5.55E-15 | 1.10E-08 | 3.86E-07 | 3.78E-07 | 3.36E-14 |
| 1 | 159452150 | 159456149 | 0.47 | 53 | 2.22E-08 | 1.15E-09 | 5.90E-10 | 4.82E-08 | 4.89E-07 | 4.46E-07 | 2.28E-09 |
| 1 | 159454150 | 159458149 | 0.31 | 49 | 4.09E-07 | 1.65E-08 | 3.97E-07 | 3.80E-07 | 9.98E-07 | 9.50E-07 | 8.56E-08 |
| 1 | 159456150 | 159460149 | 0.21 | 49 | 2.14E-04 | 4.15E-05 | 2.52E-08 | 1.59E-08 | 7.17E-07 | 6.55E-07 | 5.69E-08 |
| 1 | 159464150 | 159468149 | 0.21 | 53 | 2.37E-02 | 1.54E-03 | 2.40E-06 | 3.32E-08 | 2.60E-04 | 1.98E-04 | 1.97E-07 |
| 1 | 159468150 | 159472149 | 0.19 | 44 | 1.15E-02 | 4.07E-04 | 7.62E-09 | 2.79E-08 | 1.64E-06 | 1.35E-06 | 3.56E-08 |
| 1 | 159470150 | 159474149 | 0.29 | 46 | 8.83E-02 | 3.86E-04 | 6.39E-11 | 8.51E-08 | 2.66E-06 | 2.14E-06 | 3.83E-10 |
| 1 | 159474150 | 159478149 | 0.16 | 36 | 1.14E-06 | 4.53E-09 | 8.41E-08 | 8.43E-09 | 5.85E-06 | 5.41E-06 | 1.70E-08 |
| 1 | 159476150 | 159480149 | 0.19 | 29 | 3.42E-12 | 1.11E-14 | 7.70E-09 | 5.78E-13 | 1.36E-07 | 5.24E-08 | 6.53E-14 |
| 1 | 159478150 | 159482149 | 0.15 | 26 | 1.69E-12 | 2.11E-15 | 7.29E-11 | 7.27E-13 | 9.08E-08 | 3.91E-08 | 1.28E-14 |
| 1 | 159482150 | 159486149 | 0.21 | 44 | 3.62E-06 | 8.19E-06 | 3.20E-10 | 4.49E-10 | 1.25E-06 | 6.91E-07 | 1.12E-09 |
| 1 | 159484150 | 159488149 | 0.21 | 58 | 5.17E-06 | 2.22E-07 | 2.36E-13 | 1.09E-14 | 1.23E-06 | 6.75E-07 | 6.26E-14 |
| 1 | 159486150 | 159490149 | 0.20 | 58 | 3.91E-04 | 4.77E-06 | 1.23E-12 | 9.64E-12 | 5.14E-08 | 2.42E-08 | 6.53E-12 |
| 1 | 159488150 | 159492149 | 0.16 | 54 | 3.66E-03 | 7.66E-06 | 9.68E-14 | 4.00E-15 | 3.13E-08 | 1.67E-08 | 2.33E-14 |
| 1 | 159490150 | 159494149 | 0.27 | 48 | 4.13E-03 | 1.30E-05 | 1.02E-11 | 5.76E-10 | 2.04E-07 | 1.80E-07 | 5.99E-11 |
| 1 | 159492150 | 159496149 | 0.32 | 31 | 7.11E-04 | 3.11E-05 | 3.05E-09 | 1.24E-06 | 4.93E-07 | 5.55E-07 | 1.80E-08 |
| 1 | 159498150 | 159502149 | 0.16 | 36 | 6.55E-01 | 5.11E-02 | 4.52E-08 | 1.62E-09 | 3.09E-07 | 2.47E-07 | 9.25E-09 |
| 1 | 159500150 | 159504149 | 0.19 | 44 | 5.22E-05 | 7.89E-08 | 1.14E-12 | 1.24E-11 | 1.25E-07 | 8.63E-08 | 6.28E-12 |
| 1 | 159502150 | 159506149 | 0.24 | 54 | 9.84E-06 | 5.86E-08 | 1.33E-17 | 2.07E-11 | 1.64E-07 | 1.56E-07 | 1.11E-16 |
| 1 | 159504150 | 159508149 | 0.23 | 57 | 3.03E-02 | 8.11E-04 | 4.88E-14 | 1.05E-10 | 2.34E-08 | 2.44E-08 | 2.93E-13 |
| 1 | 159506150 | 159510149 | 0.38 | 70 | 4.66E-04 | 4.41E-06 | 4.15E-10 | 2.43E-08 | 4.47E-08 | 4.34E-08 | 2.40E-09 |
| 1 | 159512150 | 159516149 | 0.34 | 70 | 1.48E-03 | 8.58E-05 | 1.78E-09 | 6.59E-11 | 3.45E-08 | 2.26E-08 | 3.79E-10 |
| 1 | 159514150 | 159518149 | 0.24 | 44 | 5.14E-12 | 6.62E-13 | 8.44E-13 | 1.98E-12 | 1.69E-08 | 1.50E-08 | 1.77E-12 |
| 1 | 159518150 | 159522149 | 0.37 | 58 | 1.36E-04 | 1.13E-06 | 1.04E-08 | 3.70E-07 | 2.57E-08 | 2.37E-08 | 3.31E-08 |
| 1 | 159520150 | 159524149 | 0.29 | 39 | 1.17E-05 | 1.20E-08 | 2.27E-11 | 1.21E-09 | 1.96E-08 | 1.92E-08 | 1.33E-10 |
| 1 | 159534150 | 159538149 | 0.14 | 43 | 7.20E-04 | 7.47E-04 | 3.09E-10 | 2.84E-10 | 2.39E-08 | 2.07E-08 | 8.76E-10 |
| 1 | 159536150 | 159540149 | 0.46 | 65 | 1.66E-04 | 1.27E-03 | 3.76E-11 | 5.87E-12 | 8.53E-08 | 1.08E-07 | 3.05E-11 |
| 1 | 159538150 | 159542149 | 0.51 | 71 | 6.11E-06 | 1.16E-05 | 1.06E-09 | 7.51E-12 | 1.19E-07 | 1.24E-07 | 4.47E-11 |
| 1 | 159540150 | 159544149 | 0.40 | 82 | 5.10E-06 | 1.73E-09 | 4.24E-10 | 1.07E-12 | 9.11E-08 | 8.32E-08 | 6.43E-12 |
| 1 | 159542150 | 159546149 | 0.45 | 107 | 5.53E-02 | 3.41E-03 | 3.42E-08 | 1.07E-09 | 1.06E-06 | 7.46E-07 | 6.22E-09 |
| 1 | 159546150 | 159550149 | 0.17 | 45 | 3.94E-11 | 1.05E-14 | 8.54E-14 | 3.18E-18 | 9.79E-09 | 8.42E-09 | 1.11E-16 |
| 1 | 159548150 | 159552149 | 0.19 | 45 | 1.15E-07 | 4.18E-10 | 5.54E-17 | 1.38E-14 | 1.30E-08 | 1.20E-08 | 5.55E-16 |
| 1 | 159556150 | 159560149 | 0.22 | 57 | 4.39E-04 | 8.58E-03 | 1.18E-13 | 8.33E-11 | 8.87E-07 | 6.17E-07 | 7.07E-13 |
| 1 | 159558150 | 159562149 | 0.29 | 61 | 4.86E-01 | 4.83E-01 | 4.25E-09 | 3.82E-09 | 9.69E-07 | 8.39E-07 | 1.20E-08 |
| 1 | 159570150 | 159574149 | 0.30 | 82 | 6.23E-01 | 2.17E-01 | 4.42E-05 | 2.95E-04 | 2.29E-08 | 2.39E-08 | 7.01E-08 |
| 1 | 159572150 | 159576149 | 0.41 | 96 | 9.20E-01 | 9.26E-01 | 3.20E-04 | 3.05E-05 | 3.46E-08 | 3.39E-08 | 1.03E-07 |
| 1 | 159576150 | 159580149 | 0.34 | 50 | 8.99E-02 | 6.27E-01 | 2.68E-08 | 1.07E-07 | 3.74E-07 | 5.06E-07 | 1.17E-07 |
| 1 | 159580150 | 159584149 | 0.30 | 73 | 2.73E-01 | 4.90E-01 | 4.89E-10 | 2.13E-08 | 9.21E-08 | 7.75E-08 | 2.83E-09 |
| 1 | 159708150 | 159712149 | 0.30 | 56 | 1.98E-01 | 7.38E-03 | 1.50E-05 | 3.82E-12 | 7.14E-10 | 2.07E-10 | 2.24E-11 |
| 1 | 159710150 | 159714149 | 0.41 | 80 | 4.32E-02 | 8.46E-03 | 6.65E-05 | 1.90E-08 | 8.50E-10 | 2.75E-10 | 1.23E-09 |
| 1 | 159796150 | 159800149 | 0.69 | 107 | 1.95E-02 | 1.42E-01 | 4.98E-07 | 1.83E-03 | 2.77E-09 | 2.37E-09 | 7.64E-09 |
| 1 | 159798150 | 159802149 | 0.35 | 83 | 7.06E-05 | 4.62E-06 | 6.59E-13 | 1.02E-11 | 1.31E-09 | 9.13E-10 | 3.71E-12 |
| 1 | 159800150 | 159804149 | 0.16 | 61 | 3.16E-01 | 2.44E-02 | 3.56E-10 | 2.86E-10 | 3.00E-09 | 1.73E-09 | 8.31E-10 |
| 1 | 159806150 | 159810149 | 0.35 | 64 | 9.48E-01 | 3.46E-01 | 6.48E-06 | 1.29E-05 | 4.27E-08 | 3.34E-08 | 1.12E-07 |
| 1 | 159808150 | 159812149 | 0.28 | 60 | 2.12E-02 | 2.81E-03 | 6.81E-07 | 8.50E-08 | 4.89E-08 | 2.63E-08 | 8.37E-08 |
| 1 | 159810150 | 159814149 | 0.31 | 65 | 2.48E-01 | 3.69E-01 | 5.97E-06 | 5.75E-07 | 1.42E-08 | 1.04E-08 | 3.57E-08 |
| 1 | 159812150 | 159816149 | 0.44 | 70 | 8.59E-01 | 4.04E-01 | 2.43E-05 | 5.24E-08 | 1.79E-08 | 1.51E-08 | 4.26E-08 |
| 1 | 160114150 | 160118149 | 0.28 | 51 | 6.70E-04 | 2.58E-07 | 1.56E-07 | 2.22E-15 | 1.44E-13 | 4.22E-14 | 1.24E-14 |
| 1 | 160116150 | 160120149 | 0.42 | 61 | 1.26E-03 | 1.93E-05 | 5.65E-06 | 5.50E-10 | 2.24E-13 | 6.71E-14 | 3.10E-13 |
| 1 | 160614150 | 160618149 | 0.60 | 75 | 1.60E-01 | 1.75E-01 | 1.25E-07 | 5.88E-09 | 1.42E-05 | 9.12E-06 | 3.36E-08 |

| | | | | | | | | | | | |
|---|---|---|---|---|---|---|---|---|---|---|---|
| 1 | 160616150 | 160620149 | 0.55 | 73 | 1.98E-01 | 3.71E-01 | 2.41E-07 | 2.49E-08 | 1.37E-05 | 8.20E-06 | 1.35E-07 |
| 1 | 160636150 | 160640149 | 0.30 | 64 | 1.92E-02 | 1.25E-03 | 3.80E-05 | 9.22E-09 | 1.49E-05 | 5.41E-06 | 5.52E-08 |
| 1 | 161506150 | 161510149 | 0.42 | 90 | 3.31E-04 | 4.43E-03 | 3.24E-08 | 5.38E-05 | 9.83E-04 | 1.03E-03 | 1.94E-07 |
| 1 | 161508150 | 161512149 | 0.45 | 91 | 7.14E-08 | 2.57E-07 | 9.84E-10 | 3.91E-06 | 1.01E-03 | 1.13E-03 | 5.80E-09 |
| 1 | 161650150 | 161654149 | 0.24 | 71 | 8.85E-05 | 1.58E-05 | 1.42E-05 | 3.28E-08 | 1.28E-06 | 3.87E-07 | 1.76E-07 |
| 1 | 161662150 | 161666149 | 0.18 | 54 | 5.43E-03 | 2.28E-07 | 6.49E-08 | 2.55E-11 | 1.20E-06 | 3.92E-07 | 1.53E-10 |
| 1 | 161664150 | 161668149 | 0.26 | 48 | 4.42E-06 | 6.95E-11 | 3.33E-07 | 1.39E-09 | 1.05E-06 | 4.00E-07 | 3.97E-10 |
| 1 | 161670150 | 161674149 | 0.32 | 63 | 4.38E-02 | 2.40E-04 | 2.20E-06 | 3.03E-08 | 3.21E-06 | 1.26E-06 | 1.74E-07 |

cMAF = cumulative minor allele frequency
